# Nicotinamide-loaded Peptoid Nanotubes for Energy Regeneration in Acute Brain Injury

**DOI:** 10.1101/2025.03.06.641448

**Authors:** Hui Du, Hoang Trinh, Olivia C. Brandon, Renyu Zheng, Haoyu Wang, Kylie Corry, Thomas R. Wood, Chun-Long Chen, Elizabeth Nance

**Affiliations:** Department of Chemical Engineering, University of Washington, Seattle WA 98195, USA; Physical Sciences Division, Pacific Northwest National Laboratory, Richland, WA 99352, USA; Division of Neonatology, Department of Pediatrics, University of Washington, Seattle, WA 98195, USA; Department of Bioengineering, University of Washington, Seattle, WA 98195, USA; Department of Radiology, University of Washington, Seattle, WA 98195, USA

**Keywords:** ATP, cell metabolism, peptoid, nanotube, drug delivery, microglia, energy regeneration

## Abstract

Acute brain injuries such as perinatal asphyxia, stroke, and traumatic brain injury result in ischemia, oxidative stress, excitotoxicity, and inflammation, leading to a depletion of ATP, the brain’s cellular energy store. Nicotinamide adenine dinucleotide (NAD+), a key regulator of cellular homeostasis, is crucial for energy regeneration and DNA repair in post-injury recovery. However, the therapeutic benefits of NAD+ and its precursors, such as nicotinamide (NAM), are limited by the complexity of their metabolic pathways and challenges in effective cell-specific intracellular delivery. Therefore, cellular delivery strategies are needed to capture the potential of an NAD+-regenerating therapeutic approach. In this study, we introduce a nanopeptoid delivery strategy to replenish cellular redox state and energy production in the acutely injured brain. By self-assembling peptoids into tubular structures, we created biocompatible NAM-conjugated peptoid nanotubes (NAM-PNTs) that vary in tubular length. NAM-PNTs demonstrated significant therapeutic benefits by enhancing cell viability and replenishing intracellular ATP levels within 24 hours of treatment in oxygen-glucose deprived (OGD) BV-2 cells. In organotypic brain slices, NAM-PNT treatment promoted glial proliferation, reduced pro-inflammatory cytokine levels, and increased anti-inflammatory cytokines after OGD, an ex vivo model of hypoxia-ischemia. A single systemic dose of NAM-PNTs also reduced brain tissue loss and improved neuropathology after hypoxia-ischemia in term-equivalent rats. These findings highlight the strong therapeutic potential of NAM-PNTs for cell-specific targeted delivery and energy restoration in the acutely injured neonatal brain. In the neonatal brain injury field, this is the first demonstration of a novel nanoparticle platform development from first principles design and synthesis to in vitro screening and then demonstration of efficacy in vivo.

## Main

The brain is the body’s most metabolically active tissue and depends on the continuous delivery of nutrients and energy sources, including glucose and oxygen, supplied by cerebral blood flow. Through cellular respiration including glycolysis, the tricarboxylic acid (TCA) cycle, and oxidative phosphorylation, glucose is consumed to generate adenosine triphosphate (ATP), the primary cel^1^lular energy source.^2^ Nicotinamide adenine dinucleotide (NAD+) acts as an essential coenzyme in ATP production through its ability to accept hydride equivalents, forming NADH that provides electrons to the electron transport chain (ETC) to generate most ATP.^3^

Both ATP and NAD+ levels can be affected by various factors including age and health status.^4,5^ ATP and NAD+ levels in the brain are also impacted by acute injuries such as stroke, traumatic brain injury (TBI), and perinatal asphyxia, which lead to ischemia, excitotoxicity, and inflammation.^6^ These pathological processes result in mitochondrial dysfunction that reduces glycolysis and increases oxidative stress that drives DNA damage.^4^ DNA damage triggers cell death through activation of the poly(ADP-ribose) polymerase-1 (PARP-1) cell death pathway, which consumes NAD+ to repair damaged DNA. Rapid consumption of NAD+ further diminishes cellular ATP production, causing energy depletion and cell death.^7,8^ One potential way to mitigate energy depletion after injury is to directly deliver NAD+ to injured cells. For example, in a rat model of transient focal brain ischemia, intranasal administration of NAD+ was shown to decrease brain damage.^9^ Supplementation of NAD+ has also been shown to ameliorate neuropathological defects and markedly extend lifespan by promoting DNA repair and improving mitochondrial function in models of Ataxia Telangiectasia and Alzheimer’s disease (AD), decreasing neurodegenerative processes.^10–13^ Nicotinamide adenine dinucleotide (NMN), a precursor of NAD+, has been shown to have neuroprotective effect in hypoxia–ischemia^14^ and can also prevent the increase in PAR formation and NAD+ catabolism, reducing brain injury.^15^

Despite the neuroprotective potential of restoring brain NAD+ levels after injury, several issues remain to be addressed. In many studies, high and frequent doses of NAD+ and NMN are required to see benefit. In addition, NAD+ cannot pass through the cellular membrane,^16^ and the cell-specific targeting of NAD+ is limited. Exogenous NAD+ is likely to be metabolized by extracellular nucleotidases in the bloodstream, and the conversion rate of precursors like nicotinamide (NAM), NMN, and nicotinamide riboside (NR) to NAD+ in the brain is not fully understood.^17^ A recent study suggested that in the brain, NAD+ is more likely to be made by NAM-derived NMN available locally to brain cells, rather than from NR or NMN given via intravenous injection or oral delivery.^17^ Delivery vehicles could be used to reduce dose and dosing frequency, as well as increase cell specificity and protection from enzymatic degradation, in order to facilitate the import of NAD+ or NAD+ precursors into damaged brain cells.

Nanomaterials are promising delivery vehicles for intracellular drug delivery of NAD+ or NAD+ precursors. In hepatocytes, quantum dots (QDs) conjugated with NMN achieved a comparable treatment effect with a lower dose compared to free NMN.^18^ Lipid metal-organic framework (MOF) nanomaterials enabled NAD(H) to be delivered into the cellular compartment directly, replenishing the cellular NAD+ pool and increasing the survival rate after sepsis.^19^ However, toxicological studies using nanomaterials like QDs and MOFs have shown potential acute and chronic hazards to humans.^1,20^ In addition, there are no current studies on the use of biocompatible nanomaterials for the delivery of NAD+ or its precursors to specific cells in the brain.

In this study, we utilize nanotubes assembled from sequence-defined peptoids, or poly-N-substituted glycines, covalently attached with NAM to replenish NAD+ levels in the acutely injured, energy-depleted brain. Peptoids are well-advanced sequence-defined synthetic polymers developed as protein mimetics possessing advantages of both synthetic polymers and biopolymers.^21–23^ Peptoids are biocompatible and highly stable,^21^ and can be cheaply and efficiently synthesized. A wide selection of commercially available amines can attain large side-chain diversity, while exhibiting protein-like molecular recognition for applications.^21,23–25^ Due to the lack of backbone hydrogen bond donors, peptoids offer the unique simplicity for controlled assembly into nanomaterials by precisely tuning peptoid-peptoid and peptoid-surface interactions through the variation of side chains.^23,25,26^ A large number of amphiphilic peptoids have been designed and synthesized for their assembly into hierarchically structured crystalline nanomaterials with a broad range of architectures,^21,23,25^ including nanotubes.^27^

Peptoid nanotubes (PNTs) are promising biosensing, bioimaging, and drug delivery platforms^27–30^ due to their high stability, tunability, surface area, and biocompatibility in cells.^30,31^ Because nanotubes with varied lengths can extravasate more readily and easily penetrate through the tissue parenchyma and cell membrane,^32–35^ herein, we synthesized NAM-conjugated peptoid nanotubes (NAM-PNTs) and NAD+-associated PNTs of different lengths and with high drug loading efficiency. We demonstrated no cytotoxicity in brain cells or organotypic whole hemisphere (OWH) brain slices. We showed that NAM-PNTs improved cellular protection compared to free drugs by replenishing cellular ATP levels and the NAD+ pool after acute injury in oxygen-glucose deprivation (OGD)-mediated injury. After acute in vivo hypoxia-ischemia (HI), NAM-PNTs significantly reduced brain area loss and down-regulated inflammatory cytokine responses. Our study demonstrates that NAM delivery via PNTs has strong therapeutic potential for cell-specific delivery and energy regeneration in the acutely injured brain.

### Formulation and characterizations of tunable and stable NAM-PNTs

Employing a previously established solid-phase submonomer synthesis method,^36,37^ we designed and synthesized tube-forming peptoids, both with and without drug and dye molecules. Illustrated in Fig. 1a, these peptoid sequences consist of three N-(2-aminoethyl) glycine (Nae) and N-(2-carboxyethyl) glycine (Nce) groups forming the polar domain, alongside six N-[(4-bromophenyl)methyl] glycine (Nbrpm) groups comprising the hydrophobic domain. Conjugation of drug, including nicotinic acid to conjugate NAM or thymine-1-acetic acid (Thy) to electrostatically associate NAD+, or and dye molecules such as dansyl (DNS), occurred at the N-terminus adjacent to the polar domain. A detailed account of the preparation, purification, and characterizations of these four peptoids (Pep-H: Nbrpm6Nce3Nae3; Pep-NAM: Nbrpm6Nce3Nae3Nc2NAM, Pep-Thy: Nbrpm6Nce3Nae3Nc2Thy, and Pep-DNS: Nbrpm6Nce3Nae3Nc2DNS) are provided in the Methods section and supporting information (Fig. S1-4). PNTs were assembled using a method similar to evaporation-induced crystallization.^30,38^ Fig. S5 display both atomic force microscopy (AFM) and scanning electron microscopy (SEM) data, indicating that nanotubes assembled from Pep-H, Pep-NAM, Pep-Thy, and Pep-DNS exhibit a similar structure. Ex-situ AFM results indicated heights of 6.53 ± 0.29 nm (Pep-H), 7.04 ± 0.32 nm (Pep-NAM), 14.93 ± 1.25 nm (Pep-Thy), and 7.78 ± 0.07 nm (Pep-DNS). X-ray diffraction (XRD) data verified that these PNTs are highly crystalline, displaying similar XRD patterns to those PNTs we previously reported (Fig. 1b).^27–30^ The peak at 1.67 nm spacing signifies the separation between two peptoid backbones aligned with the hydrophobic Nbrpm groups facing each other.^27,28,39,40^ This observed spacing of 5.71 Å is attributed to the meticulous arrangement of aromatic side chains, specifically Nbrpm6. Additionally, a 4.61 Å spacing corresponds to the alignment of lipid-like peptoid chains. Further, the peaks at 4.32 Å, 3.81 Å, and 3.35 Å suggest extensive *π*-*π* stacking interactions^41–43^. Consistent with our prior work,^27^ the hydrophobic Nbrpm groups are densely packed together and intricately positioned at the core of the nanotube wall. The polar Nae and Nce groups are located on both surfaces of the nanotube, exhibiting a packing similar to PNTs reported previously.^27^ These results demonstrate that drug and dye molecules can be precisely incorporated within crystalline nanotubes with a tunable density while maintaining a consistent tubular framework structure.

**Figure 1.**
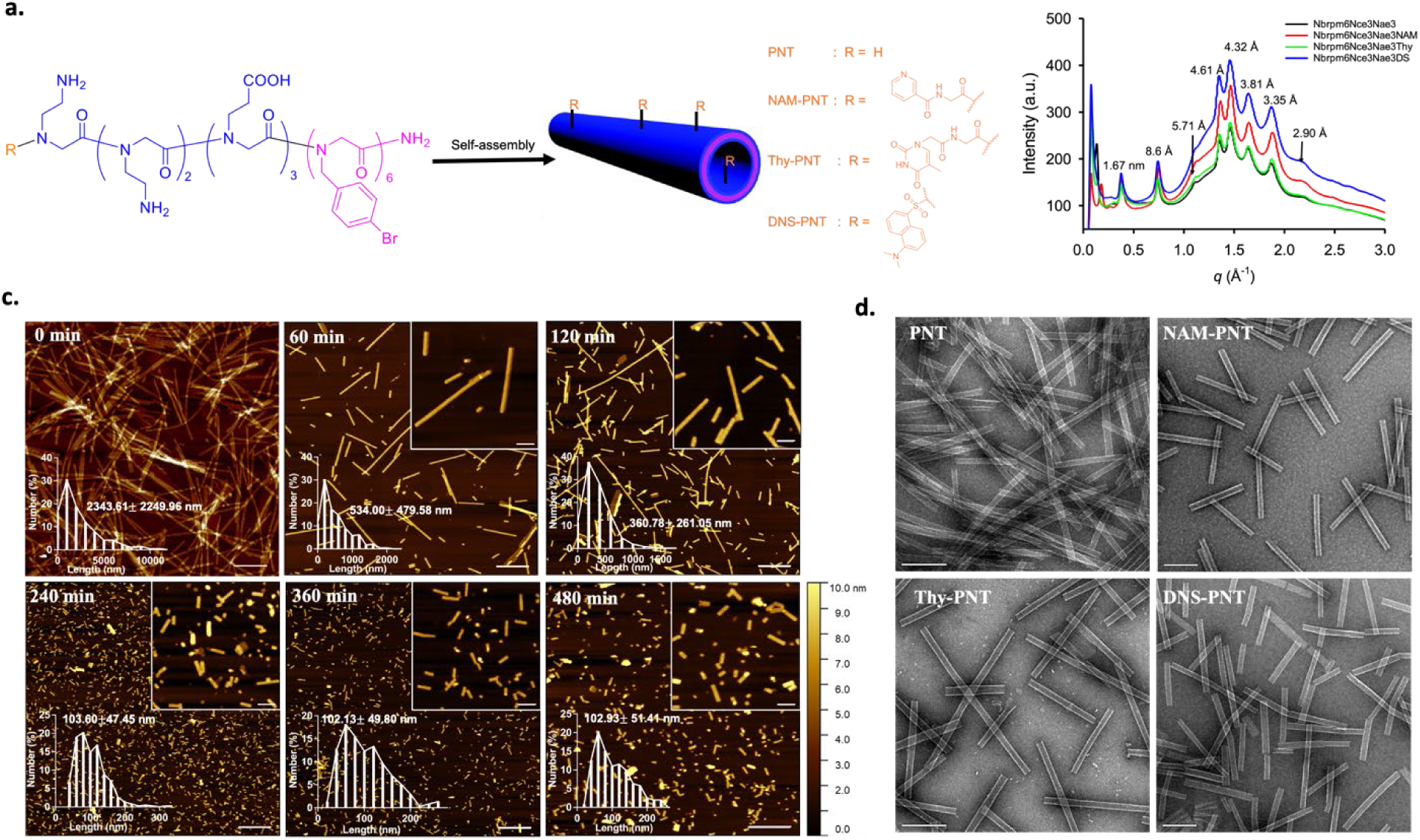
Characterizations of PNT, NAM-PNT, Thy-PNT, and DNS-PNT. a, Schematic illustration of peptoid structure and self-assembly. R indicates four different end groups in the peptoid. b, XRD data of PNT, NAM-PNT, Thy-PNT and DNS-PNT giving similar crystalline structures of all peptoids. c, AFM images showing NAM-PNTs with various lengths after 0, 60, 120, 240, 360 and 480 minutes of sonication. Scale bars, 1 µm and 200 nm (inserts). d, TEM images of PNT, NAM-PNT, Thy-PNT, and DNS-PNT, respectively. Scale bar, 200 nm.

We further showed that NAM-PNTs remained structurally unchanged in PBS over 5 days (Fig. S6). The length of PNTs can be adjusted through sonication. The length distribution of the PNT at each sonication time was quantified (Fig. S7) and the average lengths are reported in Table S1, showing reduction in length with increasing water bath sonication time (Fig. 1c). In addition to water bath sonication, probe sonication of PNTs can provide an alternative method to shorten the tube length and achieve greater monodispersity; representative TEM images of PNTs following probe sonication are shown in Fig. 1d. Sonication can cut the PNTs by inducing cavitation bubbles that release intense localized energy in the form of shockwaves and high shear forces. For NAM-PNTs, sonication did not impact NAM-conjugate stability as no NAM was released even after 360 min sonication (Fig. S8).

### NAM-PNTs show no cytotoxicity in healthy brain cells and promote energy replenishment following oxygen glucose deprivation

To confirm PNTs are biocompatible, NAM-PNTs and Thy-PNTs were tested in BV-2 cells. Dose-dependent effects on cell viability and cytotoxicity were explored with exposure to 2, 20, and 50 µg/mL PNTs. Both formulations retained high cell viability for all three dosages over 24 h of treatment (Fig. 2a, b), indicating that PNTs have no toxic effect on healthy cells, regardless of cell seeding density (Fig. S9). This result was anticipated since peptoids are known to have protein-like sequence-specific molecular recognition due to their similar structure to peptides, but confirmation in brain cells is important for the intended application. For NAM-PNTs specifically, all dosages increased cell viability. At 50 µg/mL NAM-PNTs provided a non-significant increase (11.3%) in cell viability in healthy cells compared to cells not exposed to PNTs (Fig. 2a). However, this increase was not seen in the healthy cells treated with free NAM or NAD+ (Fig. S10). In comparison, Thy-PNTs showed a non-significant decrease in cell viability with increasing Thy-PNT dose (Fig. 2b).

**Figure 2.**
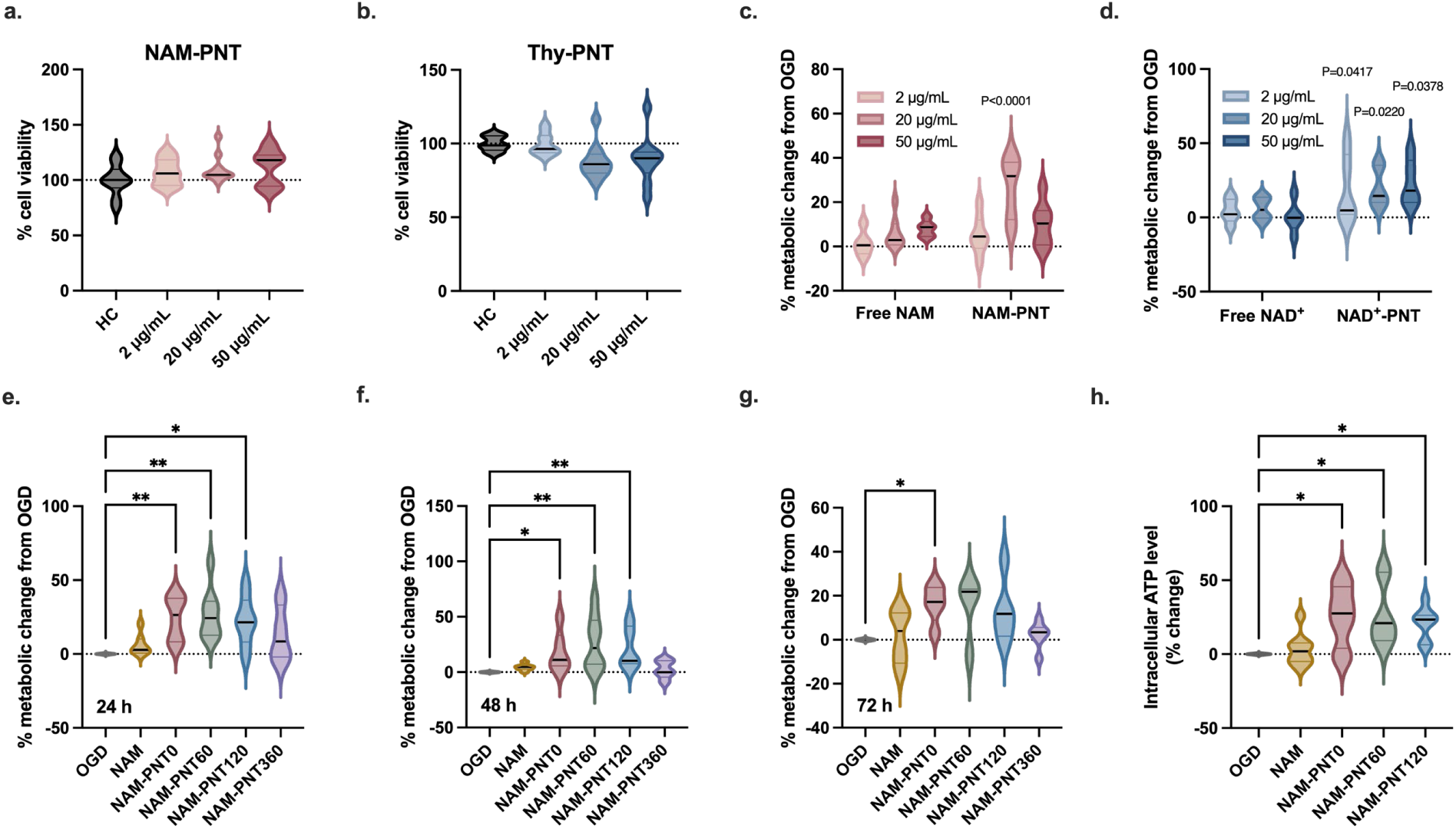
NAM-PNTs show high biocompatibility and improve cellular energy and metabolic state in the energy-depleted, acutely injured brain. a,b, MTT data showing both NAM-PNTs (a) and Thy-PNTs (b) result in negligible effects on cell viability compared to healthy control at all three dosages. (n = 10). c,d, BV-2 cells exposed to 10 minutes of OGD showed improvement in cell viability (AlamarBlue assay) after treatment with NAM-PNTs (c) and NAD+-PNTs (d) compared to NAM and NAD+ in free form. Readings were normalized to OGD controls. Statistical significance was calculated via ordinary two-way ANOVA. (n = 3-7). e, f, g, Sonication time impacts cell viability compared to non-treated OGD exposed controls after exposure to 20µg/mL of NAM-PNTs for 24 h (e), 48 h (f) and 72 h (g). Readings were normalized to OGD controls. Statistical significance was calculated via a Kruskal-Wallis test. (n = 3-6). h, Intracellular ATP level of OGD-exposed BV-2 cells with the addition of free NAM and NAM-PNTs compared to the non-treated control. Statistical significance was calculated via a Kruskal-Wallis test. All data are presented as violin plots that show the median with interquartile range.

We next sought to quantify the effect of NAM- or NAD+-PNTs after an acute injury that drives reduced glycolysis and oxidative respiration due to mitochondrial dysfunction.^44^ In this study, we applied OGD, an in vitro model of acute hypoxic-ischemic brain injury, to BV-2 cells. We first confirmed that OGD-exposed BV-2 cells showed decreased cell viability compared to healthy cells (Fig. S11). In OGD-exposed cells, the addition of NAM-PNTs increased metabolic activity, while free NAM did not (Fig. 2c). A dose-dependent effect was also observed: 20 µg/mL NAM-PNTs led to the highest increase in metabolic activity, with minimal effect at 2 µg/mL and a possible reduction in benefit at 50 µg/mL.

Delivering NAD+ directly to cells could be another method to regulate disordered metabolic states as it can decrease PARP-1-mediated cell death by maintaining glycolysis, and prevent mitochondrial depolarization and release of mitochondrial apoptosis-inducing factor.^7,45,46^ It has also been suggested that NAD+ can decrease OGD-induced neuronal death by enhancing DNA repair activity.^47,48^ However, the delivery of free NAD+ is limited, as NAD+ cannot pass through cellular membranes and has no known uptake transporters.^16^ To overcome this, NAD+ can be physically associated with Thy-PNTs via electrostatic interaction to achieve intracellular delivery. Treatment with NAD+-PNTs resulted in similar improvements in metabolic activity to those seen with NAM-PNTs (Fig. 2d), though there was no dose-dependence of effect within the doses tested. By comparison, free NAD+ had no effect.

To further investigate the efficacy of NAM-PNTs and NAD+-PNTs, we analyzed the effect of PNT length, an important physical property of a nanodelivery system that can influence cellular uptake.^49^ PNTs were cut by applying sonication. Three sonication times of 0 (PNT0), 60 (PNT60), and 120 minutes (PNT120) were applied to both NAM-PNTs and NAD+-PNTs. For all three sonication times, 20 µg/mL was determined to be the optimal dose (Fig. S12a). After 24 h of treatment, all three lengths of NAM-PNT showed comparable amounts of increase in metabolic activity compared to untreated OGD (Fig. 2e). A similar trend was measured at 48 h after NAM-PNT treatment that was sustained until 72 h (Fig. 2f, g). However, we observed that NAM-PNT360 did not improve cell viability, suggesting decreased efficacy with short PNT lengths. In healthy cells, NAM-PNT360 and NAM-PNT480 even showed an adverse effect on cell viability (Fig. S13), suggesting that much shorter PNTs may induce cell cytotoxicity, reducing the overall cell viability. Unlike NAM-PNTs, dose effect of NAD+-PNT was not clearly determined (Fig. S12) and the NAD+-PNTs showed no significant effect on metabolic change with PNT length (Fig. S14). In comparison to the chemical conjugation of NAM to PNTs, NAD+ was electrostatically associated with the PNT through molecular interactions between adenine and thymine. This association may be disrupted upon cell association or internalization, which would result in limited therapeutic efficacy observed with NAD+-PNT delivery.

Both free NAM and NAM-PNTs were able to enhance cellular ATP levels even in healthy cells over 24 h (Fig. S15a). However, free NAM showed a limited increase compared to NAM-PNTs, with a 20 µg/mL dose of NAM-PNTs giving the highest increase in cellular ATP level. By applying OGD to BV-2 cells, intracellular ATP levels decreased to 25.8% of the healthy state (Fig. S15b). Three lengths of NAM-PNTs improved ATP levels by 27.6%, 20.9% and 23.3%, respectively, after OGD energy depletion, while the free NAM did not (Fig. 2h). In an in-vitro microglial cell line, we conclude that NAM-PNTs provide a larger therapeutic potential in energy-depleted cells compared to NAD+-PNTs, but the benefit is lost the shortest PNT lengths.

### Reduced cell death via NAM-PNT treatment in OGD-exposed cultured brain slices

To capture the biocompatibility, efficacy, and cellular interaction of PNTs in a more complex model, we next evaluated NAM-PNTs in organotypic whole hemisphere (OWH) brain slices. OWH slices preserve functional relationships between neighboring cells and maintain 3D cytoarchitecture,^50^ a critical feature when trying to probe delivery into tissue. OWH brain slices also provide direct access to multiple brain regions, including the deep brain. Using OWH slices obtained from term-equivalent to human postnatal day 10 (P10) male rats, we topically applied the optimal dosage of 20 µg/mL of PNTs, as identified in our BV-2 studies, to OWH slices cultured for 4 days in vitro (DIV). We confirmed PNTs do not negatively impact cell viability in the healthy OWH slice (Fig. 3a). In fact, PNTs appear to decrease cytotoxicity compared to untreated healthy slices. When testing three different PNT lengths (NAM-PNT0, 60 and 120), we found that cytotoxicity appeared to decrease with decreasing PNT length, with PNT120 maintaining the highest cell viability, though this was not significant (p-trend=0.227).

**Figure 3.**
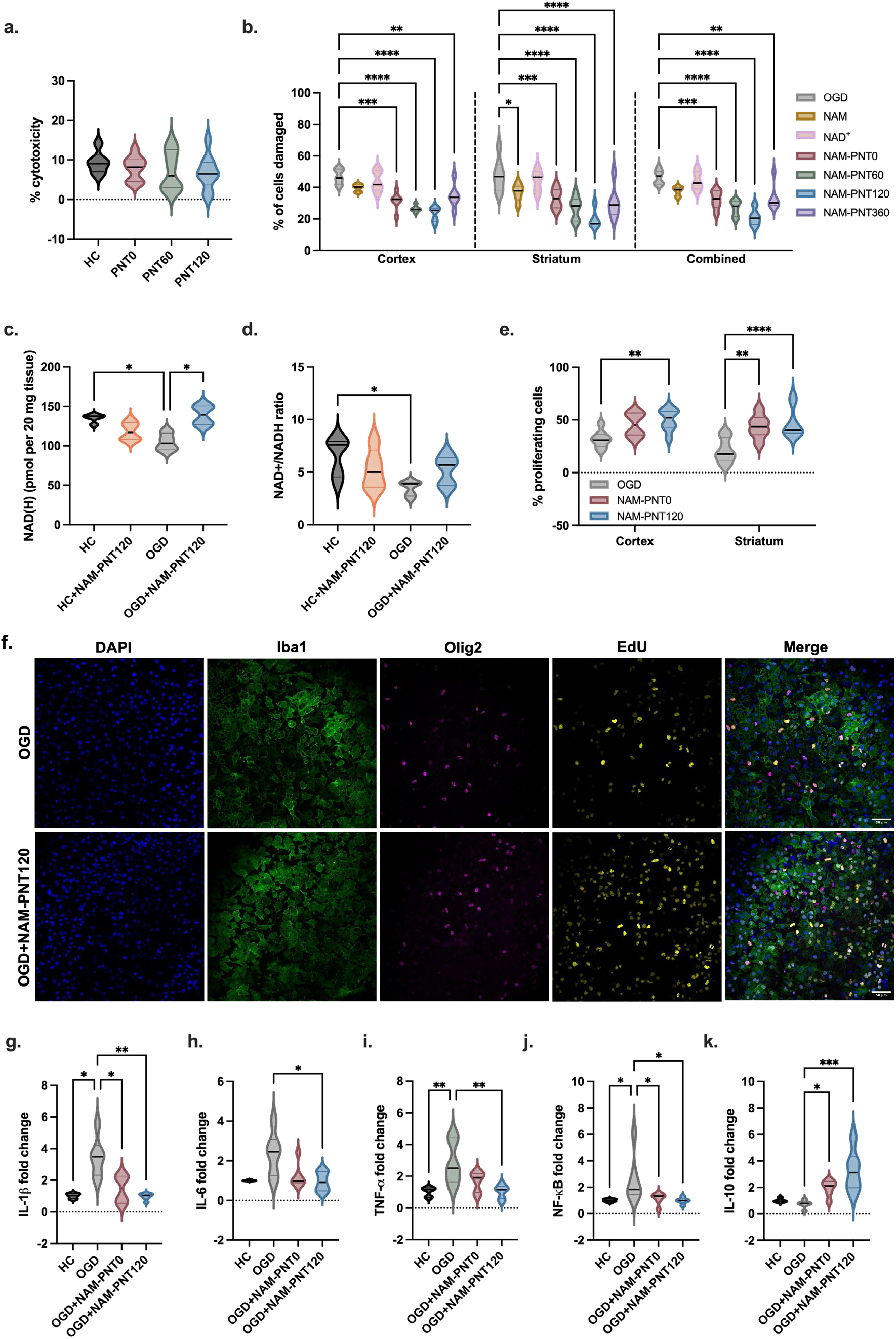
LDH, cell death and proliferation profile, and NAD level of P10 brain slices. a, LDH showing low cytotoxicity of NAM-PNTs on health slices at different sonication times. LDH release (%) values are normalized to the LDH release of acute slices immediately treated with 10% TX-100 (n = 6-9). b, NAM-PNT treatment reduces the % of damaged cells, assessed by PI quantification, in a PNT length-dependent manner. P10 profiles are given for the cortex, the striatum, and all data combined. Statistical significance was calculated via a Kruskal-Wallis test (n = 7). c, d, Intracellular NAD(H) levels (d) and NAD+/NAD(H) ratio (e) in healthy and OGD-exposed slices with or without NAM-PNT120 for 24 h. Statistical significance was calculated via a Kruskal-Wallis test (n = 5). e, Percentage of proliferating cells in P10 slices with after 30 min OGD at the cortex and the striatum. Statistical significance was calculated via ordinary two-way ANOVA (n = 6-8). f, Representative images of EdU (yellow) costaining with Iba1 (green), Olig2 (magenta), and DAPI (blue). Scale bar, 50 µm. g-k, Fold-changes of mRNA markers showing NAM-PNTs help regulating RNA expression in OGD-exposed slices with PNT treatments. Primer sequences are defined in Table S1. Statistical significance was calculated via a Kruskal-Wallis test (n = 6). All data are presented as violin plots that show the full range with lines at the median with interquartile range.

We next investigated the regional cellular response to NAM-PNT treatment in both the cortical and striatal regions of OGD-exposed OWH slices. The cortex and striatum play critical roles in normal brain function and are susceptible to hypoxic-ischemic events.^51^ In both brain regions, free NAM and free NAD+ did not improve cell viability compared to the OGD-exposed slice without treatment (Fig. 3b), as evidenced by a high number of propidium iodide (PI)-positive cells (Fig. S16). Nuclear PI accumulation is a common marker of cells undergoing necrosis.^52,53^ NAD+ is not able to permeate or transport through cellular membranes and both free NAM and NAD+ are likely metabolized by extracellular nucleosidases rather than being directly taken up into brain cells.^17^ In OGD-exposed slices, the percent of PI+ cells was reduced for all NAM-PNT treatment conditions. For both cortex and striatum, the relationship between cell death and PNT length showed cell death decreases with shorter NAM-PNTs, with NAM-PNT120 treatment resulting in the lowest percentage of damaged cells, further supporting NAM-PNT120 as a promising platform for HI treatment.

The OWH slice model requires particles to traverse the extracellular space and extracellular matrix. Prior literature has established that rod-like particles with high aspect ratio, such as single wall carbon nanotubes (SWCNTs) up to 700 nm in length, can readily diffuse in the brain cortex.^33^ Here, the therapeutic efficacy of NAM-PNTs is a trade-off with length: a higher aspect ratio contributes to cellular uptake while limiting tissue penetration rate. NAM-PNT0 have an average length of 2 µm, which are too long to penetrate through tissue, leading to a limited therapeutic efficacy. Though NAM-PNT60 showed the highest efficacy in BV-2 cells, PNT60 has a length of 500 nm with an aspect ratio of 78, which limits distribution through the intact brain tissue of the OWH slice. With increasing sonication time, the length of PNTs is reduced and the size distribution is more uniform. However, NAM-PNT360 did not show further improvement in efficacy compared to NAM-PNT120. PNTs that are too small, with a length of around 100 nm and an aspect ratio of 15, could be more rapidly cleared from the slice or have an altered extracellular and intracellular distribution that limits the effectiveness of the NAM delivery.

To assess the direct effect of NAM-PNT on NAD+ and NADH levels in healthy and OGD-exposed slices, total NAD levels with and without NAM-PNT120 were quantified. While the addition of NAM-PNT120 did not affect healthy slices, OGD-exposed slices with NAM-PNT120 had increased intracellular NAD(H) levels that were similar to healthy slices (Fig. 3c). NAM-PNT120 also improved the NAD+/NADH ratio in OGD-exposed slices (p=0.3441, Fig. 3d), but had no effect in healthy slices. Under healthy conditions, NAD+ produced from exogenous NAM is more likely to be broken down to other metabolites to maintain cellular redox state. However, when cells are NAD+ depleted, the introduction of NAM-PNT replenishes the NAD+ pool, increasing the NAD+/NADH ratio, and reducing cell death.

### NAM-PNTs drive cell proliferation, reduce pro-inflammatory responses, and restore DNA damage responses in the OGD-exposed brain

To further understand the effect of NAM-PNT120 on cellular responses to OGD, we measured cell proliferation via 5-ethynyl-2’-deoxyuridine (EdU) staining.^52,54,55^ In OGD-exposed slices, both NAM-PNT0 and NAM-PNT120 increased the number of EdU+ cells (Fig. S17). This is consistent with the PI/DAPI data as higher proliferation will result in lower relative cell death and higher metabolic activity. Regional difference in cellular proliferation was also assessed in the cortex and striatum (Fig. 3e). In both regions, NAM-PNT120 led to a higher number of EdU+ cells. In NAM-PNT120 treated OGD slices, Edu+ signal was co-localized with cellular markers of both microglia and oligodendrocytes (Fig. 3f).

OGD injury in the brain drives neuroinflammation and cell death through reduced oxidative respiration and increased oxidative stress.^56^ Before evaluating the impact of NAM-PNTs on inflammatory responses after OGD, we first applied NAM-PNT120 to healthy neonatal brain slices. We found that NAM-PNT120 resulted in no change in IL-10, TNF-α, and NF-κB, confirming the PNTs do not drive an inflammatory response (Fig. S18). In OGD-exposed slices, NAM-PNT120 was able to significantly reduce the production of IL-1β, IL-6, TNF-α, and NF-κB (Fig. 3g, h, i, j), while increasing IL-10 (Fig. 3k), thus demonstrating a strong anti-inflammatory effect. The levels of each of these markers were returned to levels comparable to the healthy control. NAM-PNT0 also reduced pro-inflammatory cytokine production, but was less effective than NAM-PNT120, confirming previous results suggesting that NAM-PNT120 provides the most effective therapeutic efficacy.

Two prototypical cytokines that are widely explored after acute brain injury are IL-6 and IL-10. IL-6 has both pro- and anti-inflammatory effects with a central role in the integrated immune defense network against infections, but in the acute phase is generally considered to be part of the pro-inflammatory injury response.^57^ IL-6 expression was increased in OGD-exposed slices and returned to the level seen in healthy slices by NAM-PNT0 and NAM-PNT120 (Fig. 3h). The anti-inflammatory cytokine IL-10 was also significantly increased compared to the OGD control group after 24 h exposure to NAM-PNTs (Fig. 3k), suggesting that PNTs play a role in alleviating inflammation in injured tissues by upregulating IL-10 mRNA expression, driving an anti-inflammatory response.

DNA damage is associated with the activation of poly(ADP-ribose) polymerase (PARP), which drives the consumption of NAD+.^58^ PARP-1 hydrolyses NAD+, providing an ADP-ribose moiety that is polymerized to poly(ADP-ribose) (PAR) in response to DNA-strand breaks.^59,60^ PAR regulates many physiological processes such as the maintenance of DNA integrity, gene expression, and cell division. Therefore, levels of PAR may act as a signature of normal DNA repair dynamics. An anti-PAR binding reagent was applied to qualitatively identify the effect of NAM-PNTs on PAR levels. Healthy slices had clear and strong PAR signals colocalizing with the nucleus, whereas the intensity of PAR was reduced in the OGD-exposed slices with a more diffuse, non-nuclear signal (Fig. S19). When PARP-1 is activated in injured cells, NAD+ can be quickly consumed to aid in DNA repair, resulting in NAD pool depletion with associated limitation of PAR production and cell death. NAM-PNT120 treatment after OGD increased the nuclear PAR signal, although some diffuse non-nuclear PAR signal remained (Fig. S19). Further mechanistic studies are needed to determine the degree to which NAM-PNT120 restores DNA repair by replenishing the NAD+ pool.

### PNTs are localized in microglia in the OGD-injured brain

To further explore PNT cellular localization, dansyl-conjugated PNTs (DNS-PNTs) were added to OWH slices. Dansyl is a fluorescent material that can be detected with high signal-to-noise even in low concentrations and is widely conjugated to peptides via N-terminal substitution.^61^ Dansyl can also be covalently attached to the primary amino groups during PNT synthesis, self-assembling into DNS-PNTs. To mirror the NAM-PNT studies, DNS-PNTs were sonicated for 0, 60, and 120 minutes and were applied to slices for 24 h. PNT lengths were comparable to those indicated in Table S1. Slices were stained with anti-Iba1 for microglia and anti-NeuN for neurons. In OWH slices, we observed large agglomerates of PNTs for DNS-PNT0, with limited colocalization in cells (Fig. 4a). With increasing sonication time, the reduced length of PNTs led to a more distributed cellular uptake in microglia (Fig. 4b). PNTs in the cells were confirmed to be intracellular and not surface-associated in microglia (Fig. 4c). PNT uptake in microglia is not surprising given the phagocytic role microglia play in maintaining a homeostatic microenvironment in the brain parenchyma.^62,63^ However, we observed an unexpected morphological change in microglia when interacting with DNS-PNT0 and DNS-PNT60. There was a significant increase in perimeter and reduction in circularity of the microglia internalizing DNS-PNT0 and DNS-PNT60 compared to DNS-PNT120 (Fig. S20). Though PNTs actively interact with microglia, the mechanism by which they are internalized remains unclear. Under healthy conditions, microglia are usually in a surveillance state, with a highly ramified phenotype. The bushy morphology with PNT0 and PNT60 may be in response to internalization of longer PNTs. The internalization of particles with high aspect ratio can distort the cell shape^64^ and the longer PNTs appear to contort microglia such that they may be internalized. Previous work has shown that PNTs that are hundreds of nanometers in length undergo endocytosis into cells;^30^ however, more mechanistic studies are required to fully elucidate PNT internalization pathways in brain cells as a function of PNT length.

**Figure 4.**
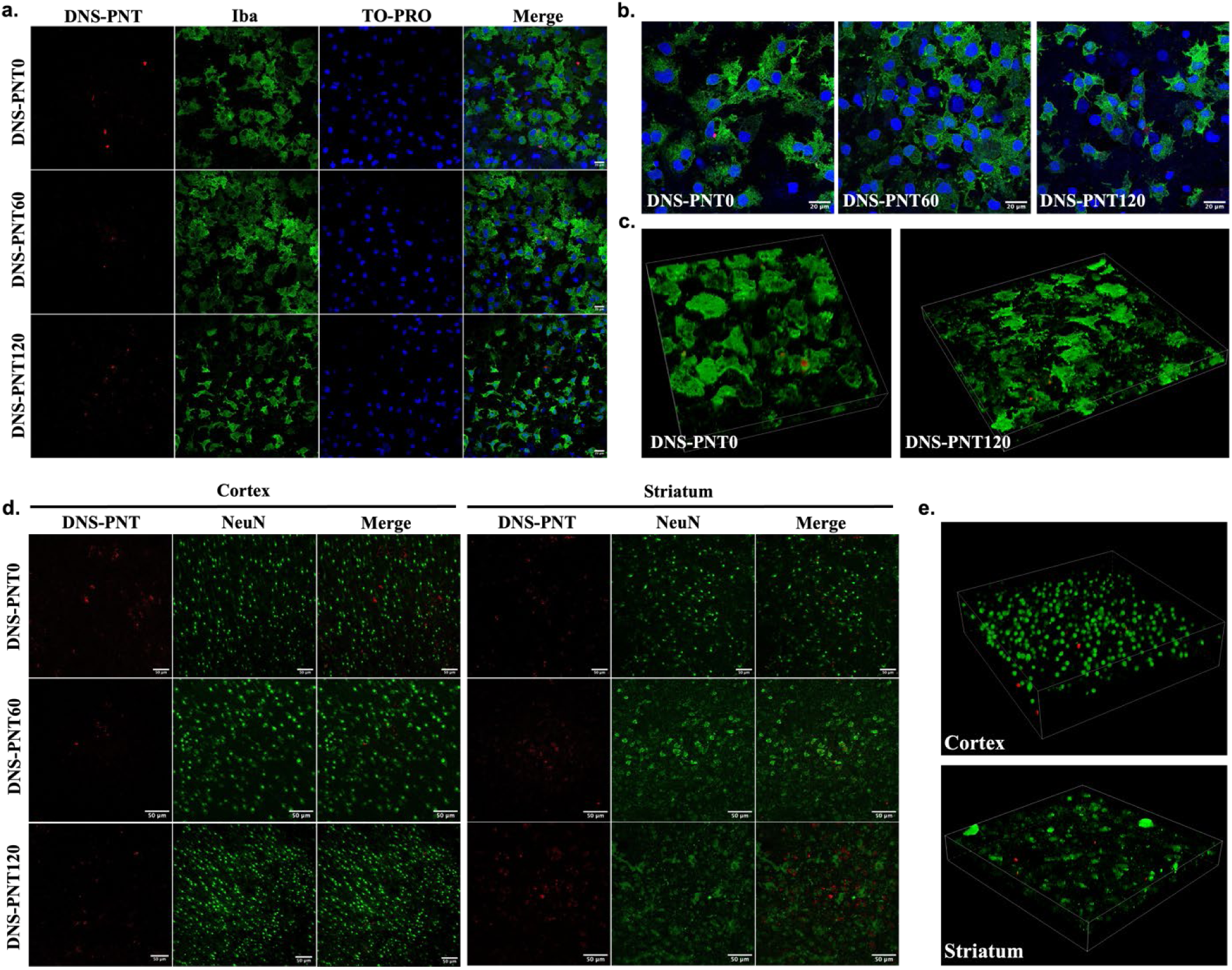
PNT localization in healthy slices. a,b, DNS-PNTs (red) localization in Iba1+ microglia (green) and ToPro-3 stained nucleus (blue) in healthy slices with magnification of 60x (a) and 100x (b). Scale bar, 50 µm. c, Z-stack of images of Iba+ microglia with DNS-PNT0 and DNS-PNT120. d, DNS-PNTs (red) localization in NeuN+ neurons (green) in healthy slices. Scale bar, 50 µm. e, Z-stack of images of NeuN+ neurons with DNS-PNT120 in cortex and striatum.

In OGD-exposed tissue, microglia have more amoeboid morphology, and PNT colocalization in microglia in OGD slices was reduced compared to healthy OWH slices, although localization was still observed for all three PNT lengths (Fig. S21). The morphology of microglia with PNT120 showed more ramified branching resembling that of a surveilling state with PNT120, compared to microglia exposed to PNT0 and 60. This may indicate the PNTs are altering the phenotype of the microglia, although more analysis of the transcriptomic state of PNT-containing microglia is needed. Neurons showed more limited interactions with PNTs, regardless of length (Fig. 4d, e), with z-stack images in both the cortex and striatum showing that DNS-PNTs had no apparent cellular interaction between the PNT and neuronal body (Fig. 4e).

### NAM-PNT treatment reduces global injury in HI-treated brain

Building on the successful optimization of NAM-PNTs in OGD-exposed OWH slices, we further evaluated the efficacy of NAM-PNTs in an in vivo hypoxia-ischemia (HI) model in P10 rats. After unilateral left carotid artery ligation and hypoxia exposure to induce unilateral HI, pups were randomized to intraperitoneal (i.p.) injection of PNTs (sonicated to length 124.24 nm, Fig. S22; 500 mg/kg), NAM-PNTs (sonicated to length 128.16 nm, Fig. S22; 25 mg/kg NAM, 500 mg/kg PNTs), free NAM (25 mg/kg), or saline (10 mL/kg) (Fig. 5a). Compared with saline, free NAM, and PNT groups, animals in the NAM-PNT lost less weight after surgery and had the highest percent (7.1%) of weight change by P13 (Fig. 5b). The median gross injury scores of all groups were at the maximum score of 4, indicating a high degree of injury; however, lower gross injury was observed with NAM-PNT treatment compared to other groups (Fig. 5c).

**Figure 5.**
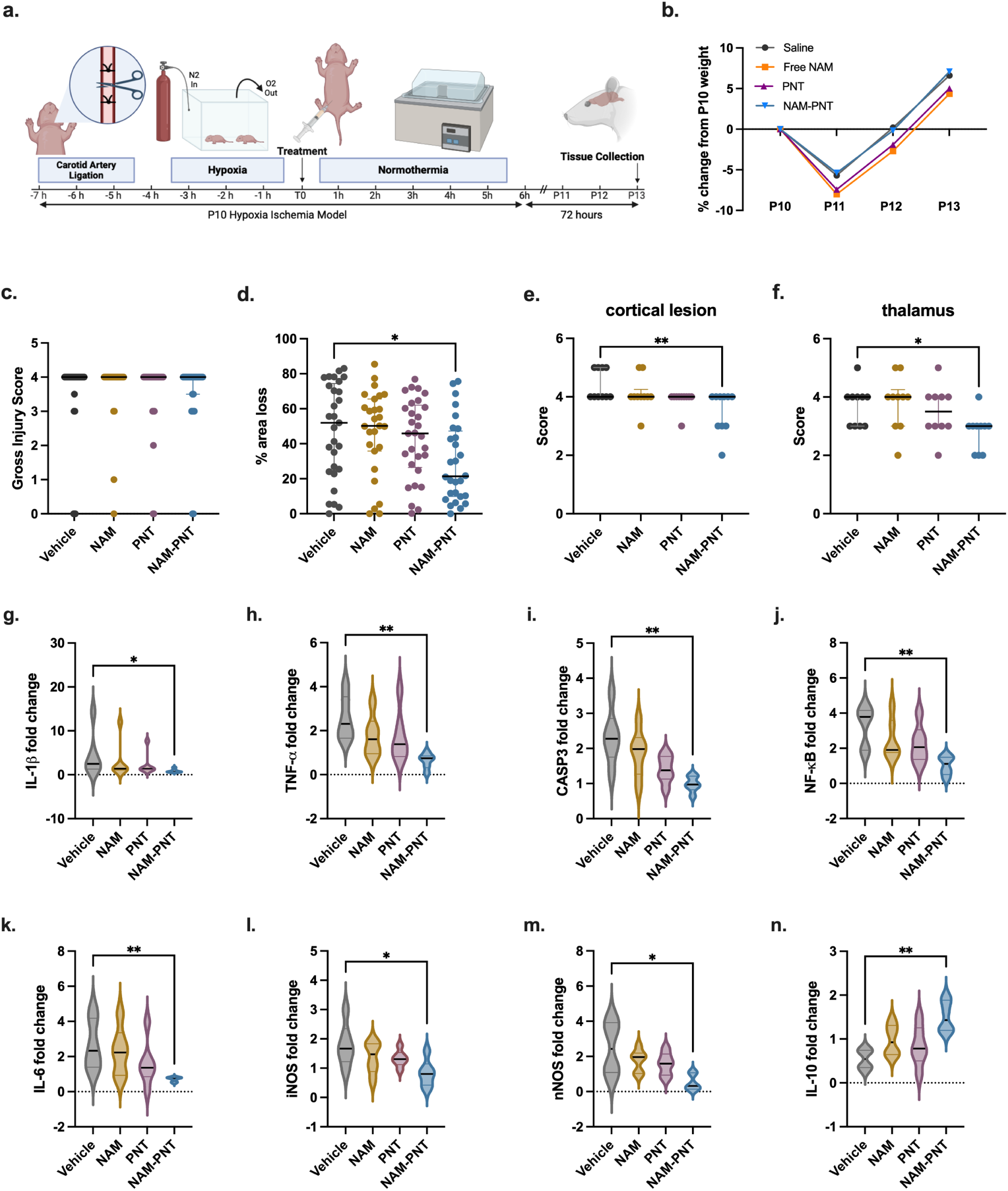
NAM-PNT effects on global brain injury and percent area loss in the HI brain. HI pups were treated 30 min after HI injury with saline (vehicle), free NAM, blank PNT, or NAM-PNT. The pups were euthanized on P13, 72 h after HI injury. All analyses were performed in a blinded manner. a, Schematic diagram of HI model. b, Weight loss compared to P10 weight after HI. c, Gross injury was assessed on a 0 (least injured) to 4 (most injured) scale for all groups: vehicle (n = 30); free NAM (n = 30); PNT (n = 31); NAM-PNT (n = 31). d, Total area of loss was calculated by assessing the percent area of tissue lost in the ipsilateral hemisphere, normalized to the contralateral hemisphere: vehicle, n = 29; free NAM, n = 27; PNT, n = 29; NAM-PNT, n = 28. Statistical significance was calculated via a Kruskal-Wallis test. e, cortical lesion score 0-5 with NAM-PNT showing significantly lower cortical lesion scoring compared to Vehicle Statistical significance was calculated via a Kruskal-Wallis test (n=10). f, thalamus neuropathology data score of 0-5 with NAM-PNT showing significantly lower lesion scoring compared to Vehicle. Statistical significance was calculated via a Kruskal-Wallis test (n=10). For plots c-e., data are presented as scatter dot plots with lines at the median with interquartile range. g-n, Fold-changes of mRNA markers compared to the contralateral hemisphere showing NAM-PNT helps regulating RNA expression in P10 rats after HI injury with PNT treatments. Statistical significance was calculated via a Kruskal-Wallis test. Data are presented as violin plots that show the full range with lines at the median with interquartile range (n = 6).

Gross injury scores are only based on the percentage of visual infarct, so further analyses assessing histopathology were conducted. Total area loss was identified by hemotoxylin & eosin (H&E) staining and was calculated by taking the ratio of injured (ipsilateral) hemisphere to the uninjured (contralateral) hemisphere.^65^ The percent area loss was significantly lower in the NAM-PNT group compared to saline (p=0.03) (Fig. 5d, Fig. S23). The NAM-PNT group had significantly lower cortical neuropathology compared to the Vehicle group (p=0.007; Fig. 5d) and less injury based on neuropathology in the thalamus (p=0.047; Fig. 5f). Presence of bilateral injury (compared to only unilateral injury) (Fig. S24a) and injury in the hippocampus (Fig. S24b) were similar between all groups.

RNA was extracted from brain tissue after HI and cytokine levels in the injured left hemisphere were compared to those in the contralateral hemisphere. The expression of proinflammatory cytokines IL-1 and TNF-α significantly decreased with NAM-PNT administration compared to the vehicle group (Fig 5.g, h). The decreased expression of cell death markers CASP3 and NF-κB were also decreased by NAM-PNT, aligning decreases in apoptosis and inflammatory activation with reduction in brain area loss (Fig 5.i, j). NAM-PNT also significantly reduced elevations in IL-6 (Fig. 5k), inducible nitric oxide synthase (iNOS), and neuronal nitric oxide synthase (nNOS) expression in the injured hemisphere compared to the contralateral hemisphere (Fig. 5l, m), as well as significantly increased expression of IL-10 (Fig. 5n). These cytokine changes further confirm that NAM-PNTs can reverse injury responses to HI in vivo.

We further explored sex-specific responses to injury and NAM-PNT, as sex can influence therapeutic efficacy in this model,^65^ and previous studies suggest that males may be more susceptible to PARP-1-related cell death.^66^ Females appeared to have higher injury (Fig. S25) as has been described previously.^65^ NAM-PNTs reduced injury by approximately 50% compared to the vehicle group, suggesting similar relative efficacy regardless of sex. Further experiments are likely to be needed to explore the role of timing and number of doses including repeat dosing over multiple days to better capture the full therapeutic potential of NAM-PNTs. However, just a single dose of NAM-PNT optimized through our in vitro and ex vivo models showed significant efficacy in vivo in the HI model.

## Conclusion

In summary, we introduce PNTs, a family of highly robust and tunable nanotubes assembled from sequence-defined synthetic polymers, as a viable brain delivery platform for NAM as an NAD+ precursor. PNTs are biocompatible without inducing an inflammatory response in the healthy brain and show localization in microglia. NAM-PNTs improved cell viability, replenished cellular energy supply and NAD(H) levels, suppressed inflammation, and reduced cell death after acute brain injury. Our findings highlight the strong therapeutic potential of NAM-PNTs for targeted delivery and energy restoration in the acutely injured neonatal brain, demonstrated through in vitro, ex vivo, and in vivo models. This study marks the first application of PNTs in neonatal disease and acute brain injury, paving the way for therapeutic strategies to drive energy regeneration in the brain. NAM-PNTs may also be applicable to other brain diseases where energy depletion is a critical driver of pathology, including other acute traumatic and ischemic brain injuries as well as chronic neurodegenerative conditions such as Parkinson’s and Alzheimer’s Disease.

## Methods

### Materials

All solvents were brought from Fisher or VWR and used without further purification. Millipore ultrapure water was employed throughout the experiments. Rink Amide resin (0.7-1.0 meq/g) and bromoacetic acid, hydroxybenzotriazole (HoBt) were purchased from Chem-Impex International, Inc. β-alanine tert-butyl ester (Nce) hydrochloride was purchased from Oakwood Chemical and was deprotected following the previous protocol before use.^36,37^ N,N’-diisopropylcarbodiimide (DIC), 4-methylpiperidine (PIP), 4-bromophenylamine (Nbrpm), tert-butyl-N-(2-aminoethyl)carbamate (Nae), and trifluoroactic acid (TFA) were purchased from Oakwood Chemical. Nicotinic acid, Fmoc-Gly-OH and thymine-1-acetic acid were purchased from Sigma-Aldrich. N,N-Diisopropylethylamine, 4-(dimethylamino)pyridine (DMAP), 1-ethyl-3-(3-dimethylaminopropyl)carbodiimide hydrochloride (EDC) and dansyl chloride (DNS) were obtained from TCI. All amine submonomers were used as received. Nicotinic acid, nicotinamide (NAM), β-nicotinamide adenine dinucleotide (NAD+), were purchased from Sigma-Aldrich. Other reagents and solvents were used as received unless otherwise stated.

### Synthesis of Functionalized Peptoids

#### Synthesis of Pep-H

The peptoids were synthesized via solid-phase synthesis, utilizing Rink resin amine as the starting material, by a previously reported method.^36,37^ Specifically, 100 mg of Rink resin amine (0.09 mmol of active NH2 groups, 1 eq.) was allowed to swell in N, N-dimethylformamide (DMF) for 10 minutes. The swollen resins were then filtered, and the Fmoc protecting groups were removed by the addition of 2 mL of a 20% (v/v) PIP/DMF solution. The resulting mixture was shaken at room temperature for 40 minutes. Following this, the resins were drained and subjected to washing with DMF through five washes of 1 mL each.

Following the deprotection step, the deprotected resins underwent an acylation reaction (Fig. 6a). This involved the use of 1.5 mL of a 0.6 M bromoacetic acid solution and 0.3 mL of a 50/50 (v/v) DIC/DMF mixture. The reaction mixture was shaken for 10 minutes at room temperature, followed by washing with DMF (5 × 1 mL). Nucleophilic displacement of bromide with the submonomers was accomplished by introducing 1.5 mL of a 0.6 M primary amine solution in N-methyl-2-pyrrolidone (NMP) and agitating the mixture for 10 minutes at room temperature. Subsequently, the solution was filtered, and the resins were washed with DMF (5 × 1 mL). The acylation and displacement reactions, employing appropriate primary amines such as Nbrpm, Nce, or Nae, were iteratively performed until the desired target peptoid sequence (termed as Nbrpm6Nce3Nae3) was achieved (Fig. 6b).

**Figure 6.**
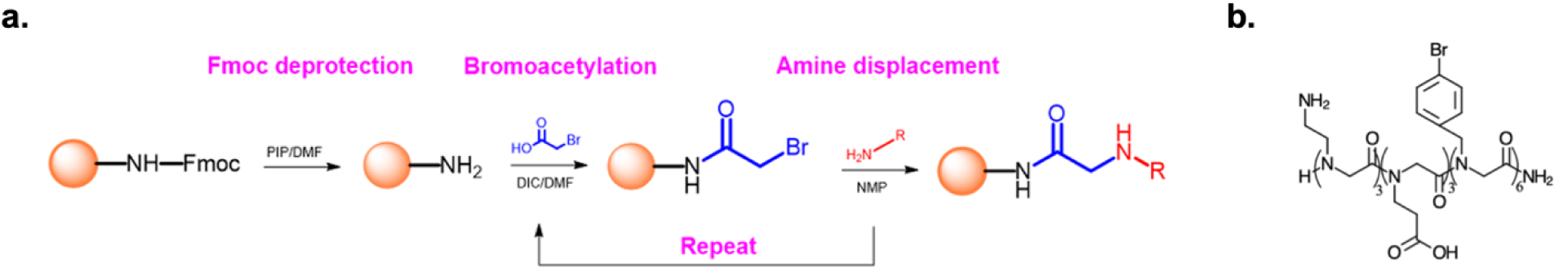
a, General procedure for submonomer synthesis of peptoids. b, Chemical structure of Nbrpm6Nce3Nae3.

#### Synthesis of Pep-NAM

To achieve NAM-containing peptoid, Nbrpm6Nce3Nae3 (0.09 mmol, 1 eq.) was first allowed to swell in DMF for 10 minutes. The swollen resins were then filtered, and Fmoc-Gly-OH (268 mg, 0.9 mmol, 10 eq.) and 2 mL of a 50/50 (v/v) DIC/DMF mixture were added. The reaction proceeded by shaking at room temperature overnight. Subsequently, the resins were drained and washed with DMF through five washes of 1 mL each. Following this, the Fmoc protecting groups were removed by adding 2 mL of a 20% (v/v) PIP/DMF solution, and the resulting mixture was shaken at room temperature for 40 minutes. The resins were then drained and washed with DMF through five washes of 1 mL each. Next, nicotinic acid (111 mg, 0.9 mmol, 10 eq.) and 2 mL of a 50/50 (v/v) DIC/DMF mixture were added, and the reaction was shaken at room temperature overnight. Afterward, the resins were drained and subjected to washing with DMF through five washes of 1 mL each.

#### Synthesis of Pep-Thy and NAD+ Association

To achieve a thymine-containing peptoid, Nbrpm6Nce3Nae3 (0.09 mmol, 1 eq.) was first allowed to swell in DMF for 10 minutes. The swollen resins were then filtered, and Fmoc-Gly-OH (268 mg, 0.9 mmol, 10 eq.) and 2 mL of a 50/50 (v/v) DIC/DMF mixture were added. The reaction proceeded by shaking at room temperature overnight. Subsequently, the resins were drained and washed with DMF through five washes of 1 mL each. Following this, the Fmoc protecting groups were removed by adding 2 mL of a 20% (v/v) PIP/DMF solution, and the resulting mixture was shaken at room temperature for 40 minutes. The resins were then drained and washed with DMF through five washes of 1 mL each. Next, thymine-1-acetic acid (166 mg, 0.9 mmol, 10 eq.), EDC (173 mg, 0.9 mmol, 10 eq.), HoBt (122 mg, 0.9 mmol, 10 eq.), DMAP (22 mg, 0.18 mmol, 2 eq.) and 2 mL of DMF were added, and the reaction was shaken at room temperature overnight. Afterward, the resins were drained and subjected to washing with DMF through five washes of 1 mL each. Before NAD+ association, 50 µL of Pep-Thy (1 µM) were sonicated for 60, 120, and 360 min for cutting into different lengths if needed. The resulting PNTs were then added into 950 mL NAD+ solution (1 mg/mL) and mixed for 30 min. The PNTs were allowed to centrifuge for 15 min at 15,000 g for sample collection. The supernatant was collected for further drug-loading analysis through liquid chromatography with an Eclipse C-18 column (4.6x150 mm, 5µm). The mobile phase was 100% DI with flow rate of 1 mL/min isocratic. NAD+ standard solutions were used to get the calibration curve each time of the experiment. The drug loading (DL) was identified as follows:

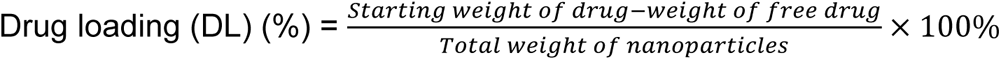

#### Synthesis of Pep-DNS

To achieve DNS-containing peptoid, Nbrpm6Nce3Nae3 (0.09 mmol, 1 eq.) was first allowed to swell in DMF for 10 minutes. Next, dansyl chloride (282 mg, 0.9 mmol, 10 eq.) and 150 µL of N,N-diisopropylethylamine were added. The reaction proceeded by shaking at room temperature overnight. Subsequently, the resins were drained and washed with DMF through five washes of 1 mL each.

#### Purification and Mass Spectrometry Analysis

The detachment of peptoids from the resin beads involved the treatment of bead resins with 3 mL of a 95/5 (v/v) trifluoroacetic acid (TFA)/H_2_O solution for 30 minutes, accompanied by agitation. The resulting solution was collected, and the TFA component was evaporated under reduced pressure at 36 °C. Subsequently, the crude peptoid product was dissolved in an 80/20 (v/v) acetonitrile/H_2_O mixture and subjected to purification through reverse-phase high-performance liquid chromatography (HPLC) using a Water 1525 system equipped with an XBridgeTM Prep C18 OBDTM column (10 µm, 19 mm × 100 mm). Purification involved employing a linear gradient of 45–55% acetonitrile in water with 0.1% TFA for Pep-NAM and Pep-THY, or 50–70% acetonitrile for other peptoids. The eluted fractions were characterized by mass spectrometry, following the methodology established in our prior work.^36^

#### Preparation of Peptoid Nanotubes and Characterization

The PNTs were prepared through the self-assembly of peptoids using the slow evaporation method. Typically, 1 µmol of peptoid was treated with 2 µL of trifluoroacetic acid (TFA) and dried under a flow of N_2_. Subsequently, 100 µL of acetonitrile was added, followed by the addition of 0.5 M NaOH. Deionized water was then added to bring the total volume to 200 µL. The samples were left at 4 °C for slow evaporation until a gel-like solution was obtained.

Atomic force microscopy (AFM) characterization: Ex situ Atomic Force Microscopy (AFM) analysis was conducted using a Bruker MultiMode 8 in ScanAsyst mode at room temperature. For sample preparation, 1.5 µL of the self-assembled peptoid was diluted 40 times with deionized water and deposited onto a mica substrate. After 5 minutes, Whatman filter paper was used to remove the solution, and then the dried mica substrate was further dried under N2 flow. The AFM probe consisted of silicon tips on silicon cantilevers (NCHV probes, k = 42 N/m, tip radius).

Scanning electron microscopy (SEM): SEM images were captured from the samples deposited on a silicon wafer using the Apreo2S scanning electron microscope by Thermo Fisher. The imaging process employed an ETD detector, an excitation voltage of 2 kV, a current of 0.2 nA, and a working distance of around 8 mm.

X-ray diffraction analysis (XRD): All characterizations were conducted following the previously described procedure.2 Powder X-ray diffraction (XRD) data were typically acquired on beamline 8.3.1, which features a 5 T single-pole superbend source with an energy range of 5– 17 keV, at the Advanced Light Source located at Lawrence Berkeley National Laboratory. XRD data were collected using a 3 × 3 CCD array (ADSC Q315r) detector with a wavelength of 1.11583 Å, and the detector was positioned 200 mm from the sample. For XRD sample preparations, peptoid assembly samples were loaded onto a Kapton mesh (MiTeGen) and dried for XRD measurements. All XRD data were analyzed using custom Python scripts.

### BV-2 Cell in vitro Model

The murine microglia cell line BV-2 was purchased from ATCC (CRL-2469) and cultured based on previous literature.^67^ Briefly, BV-2 cells were cultured in cell culture media (CCM, high-glucose, phenol red-free DMEM supplemented with 10% FBS, 1% glutamine, and 1% 100U/mL penicillin-streptomycin) at 37 ℃ in a 5% CO2 atmosphere. The passage number of BV2 used in this study was between 7 and 12. After reaching 70-80% confluency, BV-2 cells were passaged and seeded in the 96-well plate for further experiments.

### Oxygen-glucose deprivation (OGD) on BV-2 cells

OGD media consisted of 120 mM sodium chloride (NaCl, MilliporeSigma), 5 mM potassium chloride (KCl, MilliporeSigma), 2 mM calcium chloride (CaCl_2_, MilliporeSigma), 1.25 mM monosodium phosphate anhydrous (NaH_2_PO_4_, MilliporeSigma), 2 mM magnesium sulfate (MgSO_4_, MilliporeSigma), 25 mM sodium bicarbonate (NaHCO3, MilliporeSigma), and 20 mM HEPES in DI water. OGD media was sterile filtered (0.2 μm), titrated to pH 7.4 with 1 M hydrochloric acid (Thermo Fisher Scientific) or 1 M sodium hydroxide (Thermo Fisher Scientific), bubbled with nitrogen gas (Praxair, Danbury, CT, USA) for at least 10 min, and pre-warmed to 37 °C prior to use. To initiate OGD, the CCM supernatant was removed, and each well was rinsed once with 100 µL OGD media and then replenished with 100 µL fresh OGD media. The 96-well plates with OGD samples were placed in a hypoxia incubator chamber (STEMCELL Technologies, Vancouver, Canada) and placed in a 37 °C incubator. The chamber was flushed with nitrogen gas for at least 10 min then clamped shut to prevent O_2_ from entering. Cells were incubated in the chamber for another 10 min. Following OGD, wells were rinsed once with CCM and then replenished with 100 µL of fresh CCM. The cells were then returned to standard culture conditions with the desired treatments added.

### Cell cytotoxicity, viability, and ATP-level assays

BV-2 cells were seeded into a 96-well plate (1×10^4^ or 2×10^4^ per well) and cultured for two days. Subsequently, cells were treated with free NAM, NAD+, NAM-PNTs, and NAD+-PNTs (sonication times: 0, 60, 120, 240, and 360 min) after OGD for 24, 48, and 72 h. Doses were selected as 2, 20, and 50 µg/mL for PNTs, and the equivalent amount of free NAM and NAD+ was applied (0.11, 1.1, and 2.7 µg/mL). Cell cytotoxicity was identified by the standard 3-(4,5-dimethylthiazo-2-yl)-2,5-di-phenyltetrazoliumromide (MTT) assay (Thermo Fisher Scientific) on healthy cells. After 24 hours, cells were incubated with fresh media with 10 µL of 12 mM MTT solution in PBS for 4 h. Following the addition of 50 µL dimethyl sulfone (DMSO), the absorbance was recorded by a Synergy H1 multimodal microplate reader at the wavelength of 540 nm. Cell viability was measured using alamarBlueTM (aBlue) Cell Viability Reagent (Thermo Fisher Scientific). The reagent was diluted 10x in pre-warmed CCM. The existing CCM was discarded and 100 µL of aBlue CCM was added to each well. The cells were returned to the incubator and allowed to incubate for 4 h. The plate was measured directly at 560/590 nm excitation/emission fluorescence on a Synergy H1 multimodal microplate reader. Readings were normalized to healthy controls. Cellular ATP levels were tested using a luminescent ATP detection kit (catalog no. ab65348, Abcam), and performed by following the manufacturer’s protocol. Data were collected using a Synergy H1 multimodal microplate reader. For OGD studies, readings were normalized to the average value of OGD controls.

### Animal Care and Ethnic Statement for Ex vivo and In vivo Studies

All animal work was performed in accordance with the recommendations in the Guide for the Care and Use of Laboratory Animals of the National Institutes of Health (NIH). Animals were handled according to the approved Institutional Animal Care and Use Committee (IACUC) protocols (#4383-01, #4383–02, and 4484-01) of the University of Washington, Seattle, WA, USA. The University of Washington has an approved Animal Welfare Assurance (#A3464–01) on file with the NIH Office of Laboratory Animal Welfare (OLAW), is registered with the United States Department of Agriculture (USDA, certificate #91-R-0001), and is accredited by AAALAC International. Every effort was made to minimize suffering. All work was performed using Sprague-Dawley (SD) rats (virus antibody-free CD®, *Rattus norvegicus*, IGS, Charles River Laboratories, Raleigh, NC, USA) that arrived at P5-8 with a nursing dam. Dams were housed individually with their litter and allowed to acclimate to their environment. Before removal from the dam at P10, each dam and her pups were housed under standard conditions with an automatic 12 h light/dark cycle, temperature range of 20–26 °C, and access to standard chow and sterile tap water ad libitum. The pups were checked for health daily.

### Preparation of OWH Brain Slices and OGD

Healthy male SD rats were injected with an overdose of 100 μL pentobarbital (Commercial Beuthanasia D, 390 mg/mL pentobarbital, administered > 120–150 mg/kg) intraperitoneally. Once the animal was unresponsive to a toe pinch with tweezers, it was decapitated with surgical scissors. The brain was removed rapidly under aseptic conditions and submerged in ice-cold dissection media consisting of 100% HBSS (Hank’s Balanced Salt Solution, no Mg^2^+, no Ca^2+^, Thermo Fisher Scientific, Waltham, MA, USA), 1% Penicillin-Streptomycin (Thermo Fisher Scientific), and 0.64% w/v glucose (MilliporeSigma, Burlington, MA, USA). Whole brains were split into hemispheres with a sterile razor blade and sliced coronally into 300 μm-thick sections with a Mcllwain tissue chopper (Ted Pella, Inc., Redding, CA, USA). Individual slices were separated in ice-cold dissection media using sterile fine tip paint brushes and transferred onto 30-mm 0.4-μm-pore-sized cell culture inserts (hydrophilic PTFE, MilliporeSigma) before being placed in a non-treated 6-well plate (USA Scientific Inc., Ocala, FL, USA) containing 1 mL pre-warmed (37 °C) slice culture media (SCM; 50% MEM [minimum essential medium, no glutamine, no phenol red, Thermo Fisher Scientific], 21.75% HBSS [with Mg2+, with Ca2+], 25% horse serum [heat inactivated, New Zealand origin, Thermo Fisher Scientific], 1.25% HEPES [Thermo Fisher Scientific], 1% GlutaMAX Supplement, and 1% Penicillin-Streptomycin). Slices were cultured in a sterile CO2 incubator (Thermo Fisher Scientific) at 37 °C with constant humidity, 95% air and 5% CO2. All treatments were applied topically.

### OWH Sample Preparation for LDH Cytotoxicity

Two brain slices were plated per insert. At 4 DIV, 20 µg/mL of NAM-PNT0, NAM-PNT60, and NAM-PNT120 in SCM were added into slices and the SCM was collected after 24 h of incubation. To generate a positive cell death control, slices were treated for 2 h with 1% TX-100 in SCM. The supernatant was collected following TX-100 treatment. All supernatant samples were immediately stored at -80 °C. Supernatant samples were removed and thawed at room temperature to conduct LDH assays (Cayman Chemical, Ann Arbor, MI, USA). Following the manufacturer’s instructions, 100 μL of LDH reaction buffer was added to 100 μL of sample supernatant in triplicate to 96-well plates on ice, and the plates were transferred to a stir plate in a 37 °C incubator. After 30 min, absorbance at 490 nm was measured on a Synergy H1 multimode microplate reader to detect the production of colorimetric formazan. All LDH readings were normalized to an acute positive cell death control.

### Propidium Iodide (PI) Staining

After 24 h of treatments, slices were stained with 1 mL of 5 μg/mL PI (Thermo Fisher Scientific) in SCM for 45 min at standard culture conditions. The staining solution was placed underneath the insert. Slices were washed twice for 3 min each with sterile 1xPBS at room temperature, followed by a 1 h wash with 37 °C SCM at culturing conditions, and then formalin fixed for 1 h with 10% formalin phosphate buffer (Thermo Fisher Scientific). Following two final washes with room temperature 1xPBS, slices were stored covered at 4 °C until ready for use. Within 2 weeks, slices were stained for 15 min with 0.1 μg/mL DAPI (Thermo Fisher Scientific) in 1xPBS at room temperature. Slices were washed twice for 3 min each with 1xPBS prior to imaging. Two-channel 40x confocal images (oil immersion, 1.30 numerical aperture, Nikon Corporation, Minato City, Tokyo, Japan) were obtained for PI and DAPI. For every slice, five images were acquired from both the cortex and striatum. Image acquisition settings were consistent for all images and conditions. For each image, DAPI+ cell nuclei (total cells) and PI+ cell nuclei (dead cells) that were also DAPI+ were counted manually in ImageJ (NIH) after applying an Otsu threshold and fluorescent cutoff to aid in visualization. The fluorescent cutoff was kept consistent across all images. The PI+/DAPI+ cell ratio was expressed as the percentage of dead cells in an individual image.

### EdU Cell Proliferation Assay and Analysis

For proliferation analysis, Click-iT™ Plus EdU Cell Proliferation Kit for Imaging, Alexa Fluor™ 647 dye (Thermo Fisher Scientific) was used according to the manufacturer’s protocol. Briefly, EdU solution was added to live brain slices after OGD via culture media at 20 μM for 16 hours. Following fixation, an azide-conjugated dye was reacted with EdU-labeled cells in the presence of a copper catalyst for 1 h. Following a 1x PBS wash step, cellular nuclei were stained with a 0.1 μg/mL solution of DAPI in 1x PBS for 15 min prior to imaging. Three images were acquired at 40x magnification using 405 and 647 laser lines in the cortex and striatum. Each experimental group contained 3 brain slices and image acquisition settings were kept constant throughout. An ImageJ macro was used to set consistent LUT values prior to manual counting of total cells positive for DAPI and total cells positive for EdU in acquired images. The ratio of these values (EdU/DAPI) represents the percentage of proliferating cells.

### Total NAD Level and NAD+/NADH Ratio Analysis

OWH slices were treated with or without OGD at 4 DIV and treated with 20 µg/mL NAM-PNT0, and NAM-PNT120 for 24 h. Subsequently, the slices were harvested from the membrane, washed with PBS three times, and lysed using 400 µL lysis buffer. The total NAD and NAD+/NADH ratio were assessed by NAD+/NADH Quantification Kit (MAK037, Sigma-Aldrich) according to the manufacturer’s instructions.

### Immunofluorescence Staining and Imagining

Brain slices were prepared, and formalin was fixed for at least an hour. Following fixation, slices were incubated with primary antibodies for NeuN (neurons), Olig2 (oligodendrocytes), or Iba1 (microglia). On fixed slices, AlexaFluor® 488 Anti-NeuN (rabbit anti-NeuN, Abcam Cat # ab190565, Cambridge, UK) at a 1:500 dilution in 1x PBS containing 1% Triton X-100 was applied overnight at 4 °C. Following a 1x PBS wash step, slices were stored at 4 °C until they were imaged. Iba1(anti-Iba1, rabbit, FUJIFILM Wako Pure Chemical Corporation) were stained at 1:250 dilution and Olig2 (Anti-Olig2 Antibody, MilliporeSigma Cat#MABN50) stained slices were stained at 1:500 dilution in 1x PBS containing 1% Triton X-100 and 3% goat serum (MilliporeSigma, Cat#S30-100 mL) overnight at 4 °C. After washing with 1xPBS, Iba1 and Olig2 slices were stained with an AlexaFluor® 488 goat anti-rabbit secondary antibody (ThermoFisher Scientific Cat#A11034) and AlexaFluor® 555 goat anti-mouse secondary antibody (ThermoFisher Scientific Cat #A28180) at 1:500 dilution respectively in 1x PBS containing 1% Triton X-100 for 2 h. Anti-poly-ADP-ribose binding reagent (MilliporeSigma, Cat#MABE1031) followed the same staining protocol as Iba1 at a 1:1000 dilution. Either DAPI or TOPRO-3 (Thermo Fisher Scientific) was stained for those slices at a concentration of 0.1 μg/mL in 1xPBS for 30 min at room temperature. Slices were washed twice for 3 min each with 1xPBS prior to imaging.

### DNS-PNT Localization in Microglia and Neurons

After 4 DIV, 20 µg/mL DNS-PNT0, PNT60 and PNT120 were added into slices for 24 h of incubation. The slices were either stained with AlexaFluor® 488 Anti-NeuN or Iba1 (with an AlexaFluor® 488 goat anti-rabbit secondary antibody). Two-channel 40x and 60x confocal images (oil immersion, 1.30 numerical aperture, Nikon Corporation, Minato City, Tokyo, Japan) were taken to assess PNT localization. Image acquisition settings were consistent for all images and conditions.

### Reverse Transcriptase Quantitative Polymerase Chaim Reaction (RT-qPCR)

Changes in pro-inflammatory, anti-inflammatory, cell death, and oxidative stress gene expression profiles following PNT treatment were quantified using RT-qPCR. For each condition, three brain slices were cultured together on the same membrane. Slices were preserved in RNALater (Thermofisher, Waltham, MA, USA) and kept at 4 °C prior to processing to prevent RNA degradation. The RNA from homogenized brain slices were extracted with TRIzol reagent, pelleted at 15,000× g, washed several times with ultrapure DEPC water (Thermofisher, Waltham, MA, USA) and cold 70% ethanol, and the RNA final concentration was measured using a NanoDrop. cDNA was diluted to 20 ng/μL with ultrapure RNA-free water. RNA was transcribed into cDNA using Thermofisher (Waltham, MA, USA) Reverse Transcription RNA to cDNA kit. qPCR was run using the transcribed cDNA and BioRad (Hercules, CA, USA) SYBR Green Master Mix that binds to double-stranded DNA to quantitatively track the progress of DNA amplification in real time. Primers used were: Anti- or proinflammatory cytokines (IL-1β, IL-6, IL-10, NB-κB), inflammatory cytokine (TNF-α), and a housekeeping gene (GAPDH) (Table 2.1). The qPCR runs at 95 °C for 30 s, 95 °C for 5 s, and then 55 °C for 45 s for 40 cycles. The gene expression changes in OGD-conditioned and PNT treatment groups were normalized to the healthy controls to quantify fold-expression change. Results were statistically analyzed with one-way ANOVA (Kruskal–Wallis multiple comparisons tests) and normalized by the median Cq value for all samples.

**Table 1.**
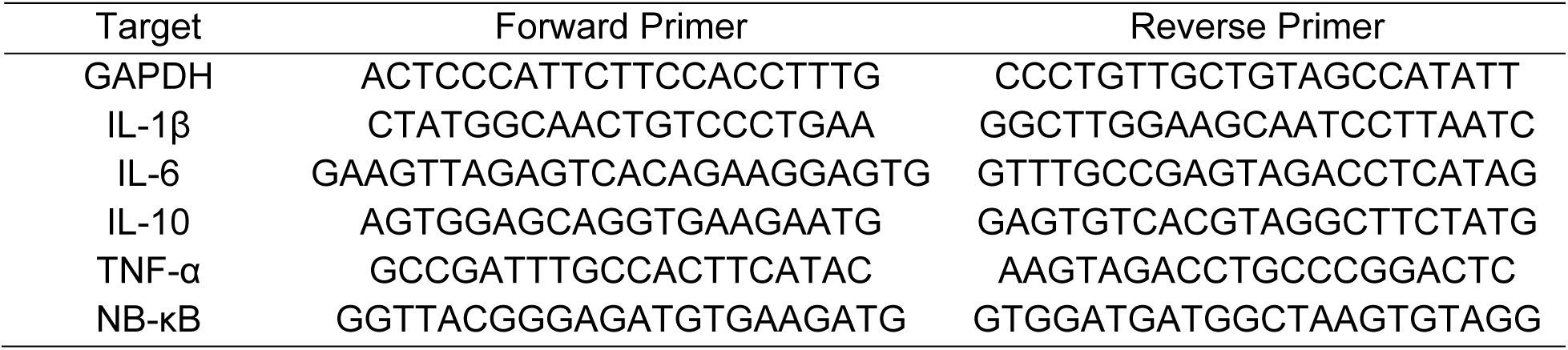
List of primers use for RNA analysis in ex vivo samples.

### Unilateral HI Brain Injury

On P10, pups were separated from their dams, weighed and sexed, and randomized to experimental groups. Buprenorphine (SQ, 0.05mg/kg) was given to all pups at least 30 minutes prior to surgery. Anesthesia with isoflurane (3% for induction, 1.5%–2.0% for maintenance) was blended with 100% O2 via a nose cone, under a dissecting microscope. A mixture of bupivacaine and lidocaine (1.5mg/kg) was injected intradermally along the incision site just prior to surgery. Under a dissecting microscope, a small (5 mm) midline neck incision was made. The left common carotid artery was identified, ligated twice with 6-0 surgical silk, and transected between the ligations. The incision was closed with tissue adhesive (Vetbond, 3M, Minnesota, USA). The pups were maintained in a temperature-controlled water bath before and after undergoing unilateral ligation of the left carotid artery to maintain normothermia. After all the animals recovered from anesthesia, they were returned to the dams for a minimum of 1 h before placement in a hypoxic chamber in a temperature-controlled water bath. Animals were sealed in the hypoxia chamber, and 8% O2 (92% N2) was administered at a rate of 2.5 L/min. During hypoxia, the temperature of the water bath was adjusted to maintain a target rectal temperature from the rectal probe of two sentinel animals at 34-35 degrees Celsius. Once the oxygen concentration within the chamber reached 8%, hypoxia was maintained for up to 180 min or 20% mortality, whichever occurred first. A research scientist continuously monitored the pups for respiration and decreased movement for the entire hypoxia duration. After the hypoxia period, the pups were then given an additional 30 min rest period with the dams. For all experiments in this study, the temperature of pups was measured immediately after the 30 min rest period. 30min after hypoxia, the dose of PNTs (500 mg/kg), saline (vehicle, 10 mL/kg), or a free drug (25 mg/kg), was administered, and the animals were then continuously monitored for 5 h in a water bath adjusted to maintain the animals at 37 degrees Celsius rectal temperature. This normothermia is critical to preventing hypothermia caused by the injury and preventing any changes in early thermoregulation that may confound the association between treatment and outcome.^68^ The animals were then returned to the dam. Animals were monitored and weighed 1-2 times daily and checked for general appearance (i.e., dehydration, abnormal posture, weight loss).

### Gross Injury and Area Loss

At 72 hours after HI, the animals received an overdose of pentobarbital before transcardiac perfusion with 1xPBS followed by 10% neutral-buffered formalin. Immediately following brain extraction, a photo of each whole brain was taken and subsequently analyzed by an individual who was blinded to group allocation. Gross brain injury in the hemisphere ipsilateral to ligation was assessed on a five-point ordinal scale (0–4) as follows: 0 = no injury, 1 = mild injury with < 25% lesion of the ipsilateral hemisphere, 2 = 25%–50% lesion, 3 = 51%–75% lesion, and 4 = ≥ 75% injury, as previously described.^69^ Whole brains were post-fixed in 10% neutral-buffered formalin for at least 48 h. Following the fixation, blocks of brain were obtained. Using external landmarks, brains were cut at approximately the level of the striatum (block 1) and at the level of the hippocampus and thalamus (block 2). The tissue samples were paraffin-embedded, cut into 5-μm sections, and stained with hematoxylin and eosin (H&E). Area loss analysis was performed as previously described.^70^ Briefly, two 5 μm sections from the slices best representing the cortex, hippocampus, basal ganglia, and thalamus were selected. Virtual slides were exported as 600 dpi images. The optical density and hemispheric area of each section were analyzed in the ImageJ software (National Institutes of Health, Bethesda, MD, USA) by another blinded individual. The average percentage area loss from the two sections (one at the level of the frontal cortex and the other at the mid hippocampal level) was calculated using the following formula: [1 − (ipsilateral area/contralateral area)] × 100.

### Histopathological Evaluation

H&E-stained slides from saline (n = 20); free NAM (n = 18); BPNT (n = 18); and NAM-PNT (n =18)-treated animals were evaluated by a board-certified veterinary pathologist blinded to the group assignment of rats. A previously reported nine-step scoring system for HIE^71^ as employed, with modifications, to grade the following regions: cerebral cortex, thalamus/midbrain, and hippocampus. Lesions in the cortex were scored semi-quantitatively using a 0–5 scale, where “0” was no detectable lesion; “1” indicated small focal or multifocal area(s) of neuronal cell loss without white matter cavitation, comprising 10% of cortex unilaterally or <5% bilaterally (3 foci or fewer), “2” indicated multifocal to coalescing lesion damage with neuronal cell loss and/or white matter cavitation, affecting 10-40% of cortex (unilateral) through all levels of cortex without cavitation; “3” indicated multifocal to coalescing lesion affecting 40-75% of cortex unilaterally or up to 33% bilaterally with or without cavitation, “4” indicated severe lesion with cavitation affecting >75% of cortex unilaterally of >50% bilaterally, and “5” indicated very severe lesions with >90% neuronal loss and extensive cavitation/collapse of cortex or striatum. Lesions of hippocampus were scored using a 0-5 scale, where “0” indicated no detectable lesion, “1” indicated rare necrotic neurons with <5% of hippocampus affected unilateral, “2” indicated multifocal mild necrotic neurons, 6-25% unilateral or up to 10% bilateral affected, “3” indicated multifocal to coalescing necrotic neurons 25-50% unilateral or 10-25% bilateral affected, “4” indicated 50-75% neurons necrotic unilateral or >25% bilateral, and “5” indicated >75% neurons necrotic. Lesions of the thalamus/midbrain were scored using a 0-5 scale, where “0” was no detectable lesion, “1” indicated small focal area to ≤3 small areas of neuronal cell loss/necrotic neurons, “2” indicated multifocal mild mf to coalescing lesions, <10%, no parenchymal loss, “3” indicated 10-25% of thalamus/midbrain affected, “4” indicated 25-50% of thalamus/midbrain affected with parenchymal loss / collapse, and “5” indicated >50% affected. In addition, an additional point was given to brains with bilateral injury.

Scores from each region were summed to yield the final score, ranging from 0 to 16. For figures, the median score from each group was calculated, and an animal representing that median score (or within 1 point of the median score) was used to show pathology in the various regions of the brain. Images of lesions captured from the digitally scanned slides were exported and plated in Adobe Photoshop Elements. Image brightness and contrast were adjusted using White Balance level and/or Auto Contrast manipulations were applied to the entire image. Original magnification and scale bars are stated.

### Statistical Analysis

Statistical analysis was performed in GraphPad Prism version 10.0.3 (GraphPad Software, San Diego, CA, USA). Graphs for the cell features compared to the control and treatment groups are displayed as the median with interquartile range and all data points are shown. Therapeutic results of free NAM, NAD+, NAM-PNT and NAD+-PNT in cells and slices were compared using Kruskal-Wallis test, expect cell viability at different doses and proliferation study in slices that uses ordinary two-way ANOVA. All p-values < 0.05 were considered statistically significant.

## Acknowledgements

The authors would like to thank Zheyu Jin for assistance in developing the initial staining immunofluorescent staining protocols and Nuo Xu for assistance in carrying in vivo administration and tissue processing procedures. The authors would like to thank Jessica M. Snyder, a board-certified pathologist, for performing blinded pathology assessment of brain sections from HI animals. The work was supported by the Jagjeet and Janice Bindra Endowed Career Development Professorship, the University of Washington Department of Chemical Engineering and the University of Washington Department of Pediatrics, Division of Neonatology. The synthesis and purification of peptoid sequences and the synthesis of PNT materials were supported by the U.S. Department of Energy (DOE), Office of Basic Energy Sciences (BES), Division of Materials Science and Engineering under an award FWP 65357 at Pacific Northwest National Laboratory (PNNL). AFM and TEM characterizations of PNTs were supported by DOE, Office of BES under award FWP 80124. Some characterizations of PNTs were carried out using facilities and equipment at the University of Washington Molecular Analysis facility, which is supported by National Nanotechnology Coordinated Infrastructure (NNCI) funded by the National Science Foundation. XRD work was conducted at the Advanced Light Source (ALS) of Lawrence Berkeley National Laboratory, which was supported by the Office of Science (No. DE-AC02-05CH11231). PNNL is multi-program national laboratory operated for Department of Energy by Battelle (No. DE-AC05-76RL01830).

## Notes

### Competing Interest Statement

The authors have declared no competing interest.

## References

1 Al Sharabati, M., Sabouni, R. & Husseini, G. A. Biomedical Applications of Metal-Organic Frameworks for Disease Diagnosis and Drug Delivery: A Review. Nanomaterials (Basel) 12 (2022). 10.3390/nano12020277

2 Dienel, G. A. Brain Glucose Metabolism: Integration of Energetics with Function. Physiol Rev 99, 949–1045 (2019). 10.1152/physrev.00062.2017

3 Blaszczyk, J. W. Energy Metabolism Decline in the Aging Brain-Pathogenesis of Neurodegenerative Disorders. Metabolites 10 (2020). 10.3390/metabo10110450

4 Verdin, E. NAD+ in aging, metabolism, and neurodegeneration. Science 350, 1208–1213 (2015). 10.1126/science.aac4854

5 Katsyuba, E., Romani, M., Hofer, D. & Auwerx, J. NAD(+) homeostasis in health and disease. Nat Metab 2, 9–31 (2020). 10.1038/s42255-019-0161-5

6 Watts, M. E., Pocock, R. & Claudianos, C. Brain Energy and Oxygen Metabolism: Emerging Role in Normal Function and Disease. Front Mol Neurosci 11, 216 (2018). 10.3389/fnmol.2018.00216

7 Alano, C. C. et al. NAD+ depletion is necessary and sufficient for poly(ADP-ribose) polymerase-1-mediated neuronal death. J Neurosci 30, 2967–2978 (2010). 10.1523/JNEUROSCI.5552-09.2010

8 Brennan, A. M. et al. NADPH oxidase is the primary source of superoxide induced by NMDA receptor activation. Nat Neurosci 12, 857–863 (2009). 10.1038/nn.2334

9 Ying, W. et al. Intranasal administration with NAD+ profoundly decreases brain injury in a rat model of transient focal ischemia. Front Biosci 12, 2728–2734 (2007). 10.2741/2267

10 Fang, E. F. et al. NAD(+) Replenishment Improves Lifespan and Healthspan in Ataxia Telangiectasia Models via Mitophagy and DNA Repair. Cell Metab 24, 566–581 (2016). 10.1016/j.cmet.2016.09.004

11 Fang, E. F. et al. Defective mitophagy in XPA via PARP-1 hyperactivation and NAD(+)/SIRT1 reduction. Cell 157, 882–896 (2014). 10.1016/j.cell.2014.03.026

12 Lautrup, S., Sinclair, D. A., Mattson, M. P. & Fang, E. F. NAD(+) in Brain Aging and Neurodegenerative Disorders. Cell Metab 30, 630–655 (2019). 10.1016/j.cmet.2019.09.001

13 Zhao, Y. et al. NAD(+) improves cognitive function and reduces neuroinflammation by ameliorating mitochondrial damage and decreasing ROS production in chronic cerebral hypoperfusion models through Sirt1/PGC-1alpha pathway. J Neuroinflammation 18, 207 (2021). 10.1186/s12974-021-02250-8

14 Kawamura, T. et al. Therapeutic Effect of Nicotinamide Mononucleotide for Hypoxic-Ischemic Brain Injury in Neonatal Mice. ASN Neuro 15, 17590914231198983 (2023). 10.1177/17590914231198983

15 Park, J. H., Long, A., Owens, K. & Kristian, T. Nicotinamide mononucleotide inhibits post-ischemic NAD(+) degradation and dramatically ameliorates brain damage following global cerebral ischemia. Neurobiol Dis 95, 102–110 (2016). 10.1016/j.nbd.2016.07.018

16 van Roermund, C. W., Elgersma, Y., Singh, N., Wanders, R. J. & Tabak, H. F. The membrane of peroxisomes in Saccharomyces cerevisiae is impermeable to NAD(H) and acetyl-CoA under in vivo conditions. EMBO J 14, 3480–3486 (1995). 10.1002/j.1460-2075.1995.tb07354.x

17 Liu, L. et al. Quantitative Analysis of NAD Synthesis-Breakdown Fluxes. Cell Metab 27, 1067–1080 e1065 (2018). 10.1016/j.cmet.2018.03.018

18 Hunt, N. J. et al. Quantum Dot Nanomedicine Formulations Dramatically Improve Pharmacological Properties and Alter Uptake Pathways of Metformin and Nicotinamide Mononucleotide in Aging Mice. ACS Nano 15, 4710–4727 (2021). 10.1021/acsnano.0c09278

19 Ye, M. et al. NAD(H)-loaded nanoparticles for efficient sepsis therapy via modulating immune and vascular homeostasis. Nat Nanotechnol 17, 880–890 (2022). 10.1038/s41565-022-01137-w

20 Hardman, R. A toxicologic review of quantum dots: toxicity depends on physicochemical and environmental factors. Environ Health Perspect 114, 165–172 (2006). 10.1289/ehp.8284

21 Sun, J. & Zuckermann, R. N. Peptoid polymers: a highly designable bioinspired material. ACS Nano 7, 4715–4732 (2013). 10.1021/nn4015714

22 Gangloff, N., Ulbricht, J., Lorson, T., Schlaad, H. & Luxenhofer, R. Peptoids and Polypeptoids at the Frontier of Supra- and Macromolecular Engineering. Chem Rev 116, 1753–1802 (2016). 10.1021/acs.chemrev.5b00201

23 Li, Z., Cai, B., Yang, W. & Chen, C.-L. Hierarchical Nanomaterials Assembled from Peptoids and Other Sequence-Defined Synthetic Polymers. Chem. Rev. 121, 14031–14087 (2021). 10.1021/acs.chemrev.1c00024

24 Yang, W., Yin, Q. & Chen, C.-L. Designing Sequence-Defined Peptoids for Biomimetic Control over Inorganic Crystallization. Chem. Mater. 33, 3047–3065 (2021). 10.1021/acs.chemmater.1c00243

25 Cai, B., Li, Z. & Chen, C.-L. Programming Amphiphilic Peptoid Oligomers for Hierarchical Assembly and Inorganic Crystallization. Acc. Chem. Res. 54, 81–91 (2021). 10.1021/acs.accounts.0c00533

26 Shao, L. et al. Hierarchical Materials from High Information Content Macromolecular Building Blocks: Construction, Dynamic Interventions, and Prediction. Chem. Rev. 122, 17397–17478 (2022). 10.1021/acs.chemrev.2c00220

27 Jin, H. et al. Designable and dynamic single-walled stiff nanotubes assembled from sequence-defined peptoids. Nat Commun 9, 270 (2018). 10.1038/s41467-017-02059-1

28 Song, Y. et al. Assembly of highly efficient aqueous light-harvesting system from sequence-defined peptoids for cytosolic microRNA detection. Nano Research 17, 788–796 (2024). 10.1007/s12274-023-6008-0

29 Cai, X. et al. Sequence-Defined Nanotubes Assembled from IR780-Conjugated Peptoids for Chemophototherapy of Malignant Glioma. Research 2021, 9861384 (2021). 10.34133/2021/9861384

30 Luo, Y. et al. Bioinspired Peptoid Nanotubes for Targeted Tumor Cell Imaging and Chemo-Photodynamic Therapy. Small 15, e1902485 (2019). 10.1002/smll.201902485

31 Song, Y. et al. Efficient Cytosolic Delivery Using Crystalline Nanoflowers Assembled from Fluorinated Peptoids. Small 14, e1803544 (2018). 10.1002/smll.201803544

32 Mitchell, M. J. et al. Engineering precision nanoparticles for drug delivery. Nat Rev Drug Discov 20, 101–124 (2021). 10.1038/s41573-020-0090-8

33 Godin, A. G. et al. Single-nanotube tracking reveals the nanoscale organization of the extracellular space in the live brain. Nat Nanotechnol 12, 238–243 (2017). 10.1038/nnano.2016.248

34 Salatin, S., Maleki Dizaj, S. & Yari Khosroushahi, A. Effect of the surface modification, size, and shape on cellular uptake of nanoparticles. Cell Biol Int 39, 881–890 (2015). 10.1002/cbin.10459

35 Ma, N. et al. Influence of nanoparticle shape, size, and surface functionalization on cellular uptake. J Nanosci Nanotechnol 13, 6485–6498 (2013). 10.1166/jnn.2013.7525

36 Wang, M. et al. Programmable two-dimensional nanocrystals assembled from POSS-containing peptoids as efficient artificial light-harvesting systems. Sci Adv 7 (2021). 10.1126/sciadv.abg1448

37 Jin, H. et al. Highly stable and self-repairing membrane-mimetic 2D nanomaterials assembled from lipid-like peptoids. Nat Commun 7, 12252 (2016). 10.1038/ncomms12252

38 Wang, M. et al. Peptoid-Based Programmable 2D Nanomaterial Sensor for Selective and Sensitive Detection of H(2)S in Live Cells. ACS Appl Bio Mater 3, 6039–6048 (2020). 10.1021/acsabm.0c00657

39 Jian, T. et al. Highly stable and tunable peptoid/hemin enzymatic mimetics with natural peroxidase-like activities. Nat. Commun. 13, 3025 (2022). 10.1038/s41467-022-30285-9

40 Li, Z. et al. Amphiphilic Peptoid-Directed Assembly of Oligoanilines into Highly Crystalline Conducting Nanotubes. Macromol. Rapid Commun. 43, 2100639 (2022). 10.1002/marc.202100639

41 Cai, Y.-P. et al. Self-assembly of silver(I) polymers with single strand double-helical structures containing the ligand O,O′-bis(8-quinolyl)-1,8-dioxaoctane. *Journal of the Chemical Society, Dalton Transactions*, 2429-2434 (2001). 10.1039/B102525M

42 Chen, C. L., Tan, H. Y., Yao, J. H., Wan, Y. Q. & Su, C. Y. Disilver(I) rectangular-shaped metallacycles: X-ray crystal structure and dynamic behavior in solution. Inorg. Chem. 44, 8510–8520 (2005). 10.1021/ic050673o

43 Chen, C. L. & Beatty, A. M. Guest inclusion and structural dynamics in 2-D hydrogen-bonded metal-organic frameworks. J. Am. Chem. Soc. 130, 17222–17223 (2008). 10.1021/ja806180z

44 Bhatti, J. S., Bhatti, G. K. & Reddy, P. H. Mitochondrial dysfunction and oxidative stress in metabolic disorders - A step towards mitochondria based therapeutic strategies. Biochim Biophys Acta Mol Basis Dis 1863, 1066–1077 (2017). 10.1016/j.bbadis.2016.11.010

45 Ying, W., Garnier, P. & Swanson, R. A. NAD+ repletion prevents PARP-1-induced glycolytic blockade and cell death in cultured mouse astrocytes. Biochem Biophys Res Commun 308, 809–813 (2003). 10.1016/s0006-291x(03)01483-9

46 Alano, C. C., Ying, W. & Swanson, R. A. Poly(ADP-ribose) polymerase-1-mediated cell death in astrocytes requires NAD+ depletion and mitochondrial permeability transition. J Biol Chem 279, 18895–18902 (2004). 10.1074/jbc.M313329200

47 Khoury, N., Koronowski, K. B., Young, J. I. & Perez-Pinzon, M. A. The NAD(+)-Dependent Family of Sirtuins in Cerebral Ischemia and Preconditioning. Antioxid Redox Signal 28, 691–710 (2018). 10.1089/ars.2017.7258

48 Wang, S. et al. Cellular NAD replenishment confers marked neuroprotection against ischemic cell death: role of enhanced DNA repair. Stroke 39, 2587–2595 (2008). 10.1161/STROKEAHA.107.509158

49 Chithrani, B. D., Ghazani, A. A. & Chan, W. C. Determining the size and shape dependence of gold nanoparticle uptake into mammalian cells. Nano Lett 6, 662–668 (2006). 10.1021/nl052396o

50 Humpel, C. Organotypic brain slice cultures: A review. Neuroscience 305, 86–98 (2015). 10.1016/j.neuroscience.2015.07.086

51 Wise-Faberowski, L., Robinson, P. N., Rich, S. & Warner, D. S. Oxygen and glucose deprivation in an organotypic hippocampal slice model of the developing rat brain: the effects on N-methyl-D-aspartate subunit composition. Anesth Analg 109, 205–210 (2009). 10.1213/ane.0b013e3181a27e37

52 Ferlini, C., Biselli, R., Scambia, G. & Fattorossi, A. Probing chromatin structure in the early phases of apoptosis. Cell Prolif 29, 427–436 (1996). 10.1111/j.1365-2184.1996.tb00985.x

53 Singh, N. P. A simple method for accurate estimation of apoptotic cells. Exp Cell Res 256, 328–337 (2000). 10.1006/excr.2000.4810

54 Salic, A. & Mitchison, T. J. A chemical method for fast and sensitive detection of DNA synthesis in vivo. Proc Natl Acad Sci U S A 105, 2415–2420 (2008). 10.1073/pnas.0712168105

55 Cappella, P., Gasparri, F., Pulici, M. & Moll, J. A novel method based on click chemistry, which overcomes limitations of cell cycle analysis by classical determination of BrdU incorporation, allowing multiplex antibody staining. Cytometry A 73, 626–636 (2008). 10.1002/cyto.a.20582

56 Teleanu, D. M. et al. An Overview of Oxidative Stress, Neuroinflammation, and Neurodegenerative Diseases. Int J Mol Sci 23 (2022). 10.3390/ijms23115938

57 Tanaka, T., Narazaki, M. & Kishimoto, T. IL-6 in inflammation, immunity, and disease. Cold Spring Harb Perspect Biol 6, a016295 (2014). 10.1101/cshperspect.a016295

58 Murata, M. M. et al. NAD+ consumption by PARP1 in response to DNA damage triggers metabolic shift critical for damaged cell survival. Mol Biol Cell 30, 2584–2597 (2019). 10.1091/mbc.E18-10-0650

59 D’Amours, D., Sallmann, F. R., Dixit, V. M. & Poirier, G. G. Gain-of-function of poly(ADP-ribose) polymerase-1 upon cleavage by apoptotic proteases: implications for apoptosis. J Cell Sci 114, 3771–3778 (2001). 10.1242/jcs.114.20.3771

60 Ray Chaudhuri, A. & Nussenzweig, A. The multifaceted roles of PARP1 in DNA repair and chromatin remodelling. Nat Rev Mol Cell Biol 18, 610–621 (2017). 10.1038/nrm.2017.53

61 Weber, G. Polarization of the fluorescence of macromolecules. II. Fluorescent conjugates of ovalbumin and bovine serum albumin. Biochem J 51, 155–167 (1952). 10.1042/bj0510155

62 Butovsky, O. & Weiner, H. L. Microglial signatures and their role in health and disease. Nat Rev Neurosci 19, 622–635 (2018). 10.1038/s41583-018-0057-5

63 Li, Q. & Barres, B. A. Microglia and macrophages in brain homeostasis and disease. Nat Rev Immunol 18, 225–242 (2018). 10.1038/nri.2017.125

64 Champion, J. A. & Mitragotri, S. Shape Induced Inhibition of Phagocytosis of Polymer Particles. Pharmaceutical Research 26, 244–249 (2009). 10.1007/s11095-008-9626-z

65 Wood, T. R. et al. Variability and sex-dependence of hypothermic neuroprotection in a rat model of neonatal hypoxic-ischaemic brain injury: a single laboratory meta-analysis. Sci Rep 10, 10833 (2020). 10.1038/s41598-020-67532-2

66 Hagberg, H. et al. PARP-1 gene disruption in mice preferentially protects males from perinatal brain injury. J Neurochem 90, 1068–1075 (2004). 10.1111/j.1471-4159.2004.02547.x

67 Zhang, M. et al. Quantum Dot Cellular Uptake and Toxicity in the Developing Brain: Implications for Use as Imaging Probes. Nanoscale Adv 1, 3424–3442 (2019). 10.1039/C9NA00334G

68 Galinsky, R. et al. In the Era of Therapeutic Hypothermia, How Well Do Studies of Perinatal Neuroprotection Control Temperature? Dev Neurosci 39, 7–22 (2017). 10.1159/000452859

69 Juul, S. E. et al. Microarray analysis of high-dose recombinant erythropoietin treatment of unilateral brain injury in neonatal mouse hippocampus. Pediatr Res 65, 485–492 (2009). 10.1203/PDR.0b013e31819d90c8

70 Sabir, H., Scull-Brown, E., Liu, X. & Thoresen, M. Immediate hypothermia is not neuroprotective after severe hypoxia-ischemia and is deleterious when delayed by 12 hours in neonatal rats. Stroke 43, 3364–3370 (2012). 10.1161/STROKEAHA.112.674481

71 Thoresen, M., Bagenholm, R., Loberg, E. M., Apricena, F. & Kjellmer, I. Posthypoxic cooling of neonatal rats provides protection against brain injury. Arch Dis Child Fetal Neonatal Ed 74, F3–9 (1996). 10.1136/fn.74.1.f3

