## Supplemental Figures 1-25 and Table S1 for "Nicotinamide-loaded Peptoid Nanotubes for Energy Regeneration in Acute Brain Injury"

Supplemental Information

Hui Du[1], Hoang Trinh[2], Olivia C. Brandon[3], Renyu Zheng[1,2], Haoyu Wang[1], Kylie Corry[3], Thomas R. Wood[3], Chun-Long Chen[1,2], and Elizabeth Nance[1,4,5]*

[1]Department of Chemical Engineering, University of Washington, Seattle WA 98195, USA
[2]Physical Sciences Division, Pacific Northwest National Laboratory, Richland, WA 99352, USA.
[3]Division of Neonatology, Department of Pediatrics, University of Washington, Seattle, WA 98195, USA
[4]Department of Bioengineering, University of Washington, Seattle, WA 98195, USA
[5]Department of Radiology, University of Washington, Seattle, WA 98195, USA

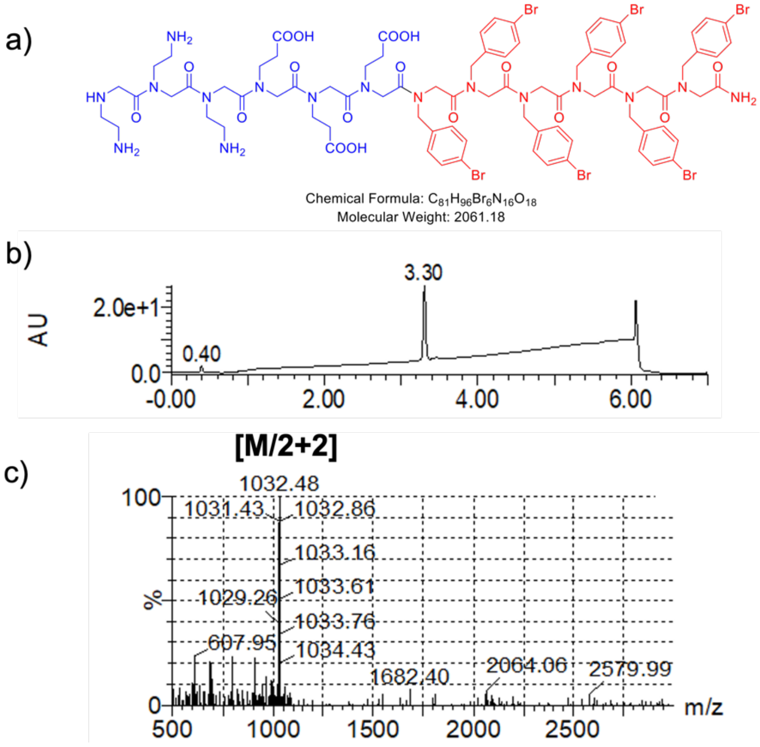


Figure S1. LC-MS analysis of PNT. a) Chemical structure of PNT. b) LC profile of PNT run under gradient of acetonitrile from 5 -95 %. c) ESI mass analysis of PNT.

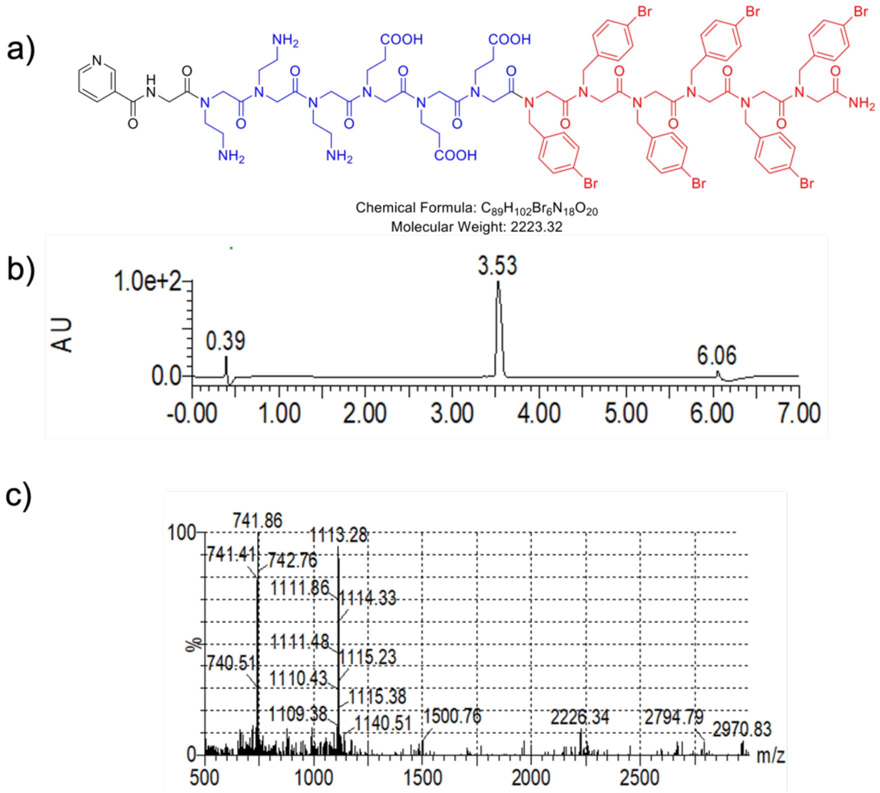


Figure S2. LC-MS analysis of NAM-PNT. a) Chemical structure of NAM-PNT. b) LC profile of NAM-PNT run under gradient of acetonitrile from 5 -95 %. c) ESI mass analysis of NAM-PNT.

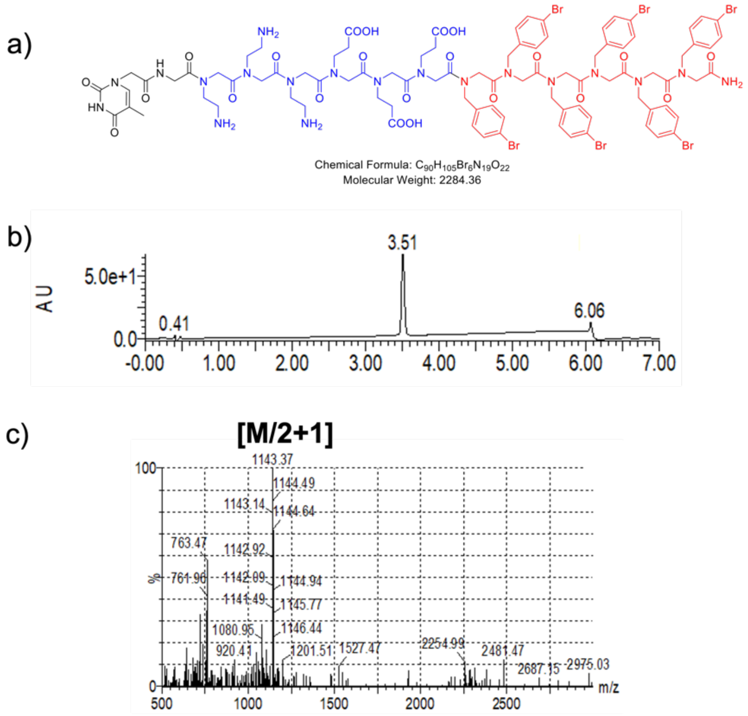


Figure S3. LC-MS analysis of Thy-PNT. a) Chemical structure of Thy-PNT. b) LC profile of Thy-PNT run under gradient of acetonitrile from 5 -95 %. c) ESI mass analysis of Thy-PNT.

d)

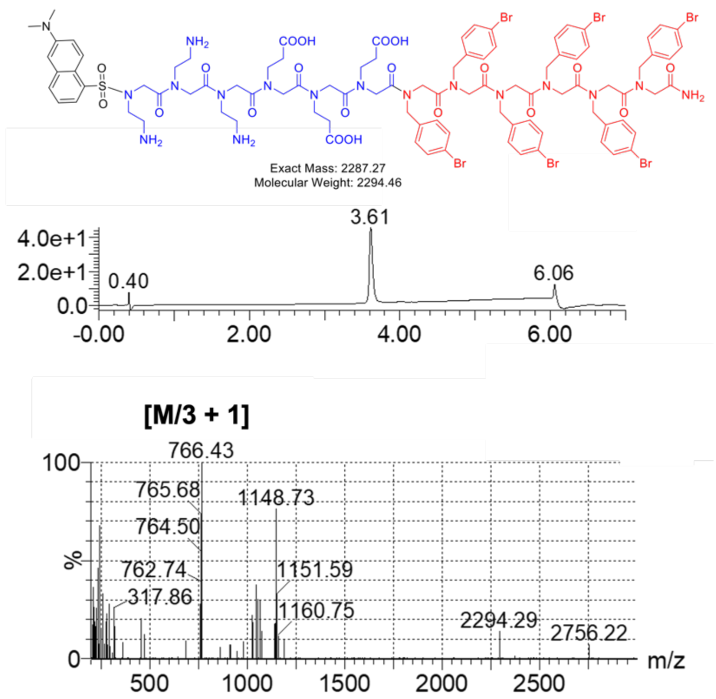


Figure S4. LC-MS analysis of DS-PNT. a) Chemical structure of DS-PNT. b) LC profile of DS-PNT run under gradient of acetonitrile from 5 -95 %. c) ESI mass analysis of DS-PNT.

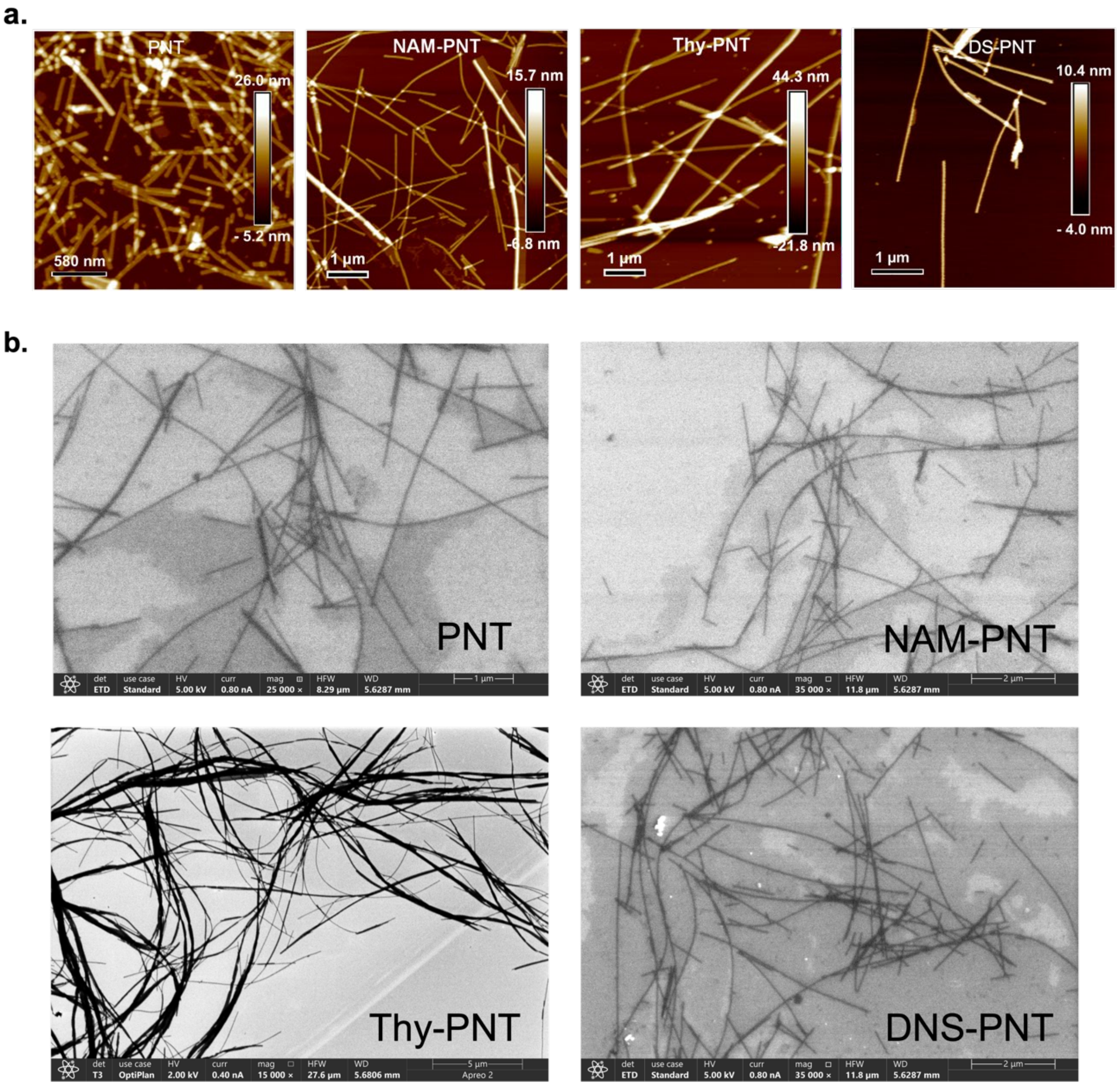


Figure S5. a) AFM images of PNT, NAM-PNT, Thy-PNT, and DNS-PNT, respectively. b) SEM images of PNT, NAM-PNT, Thy-PNT and DNS-PNT.

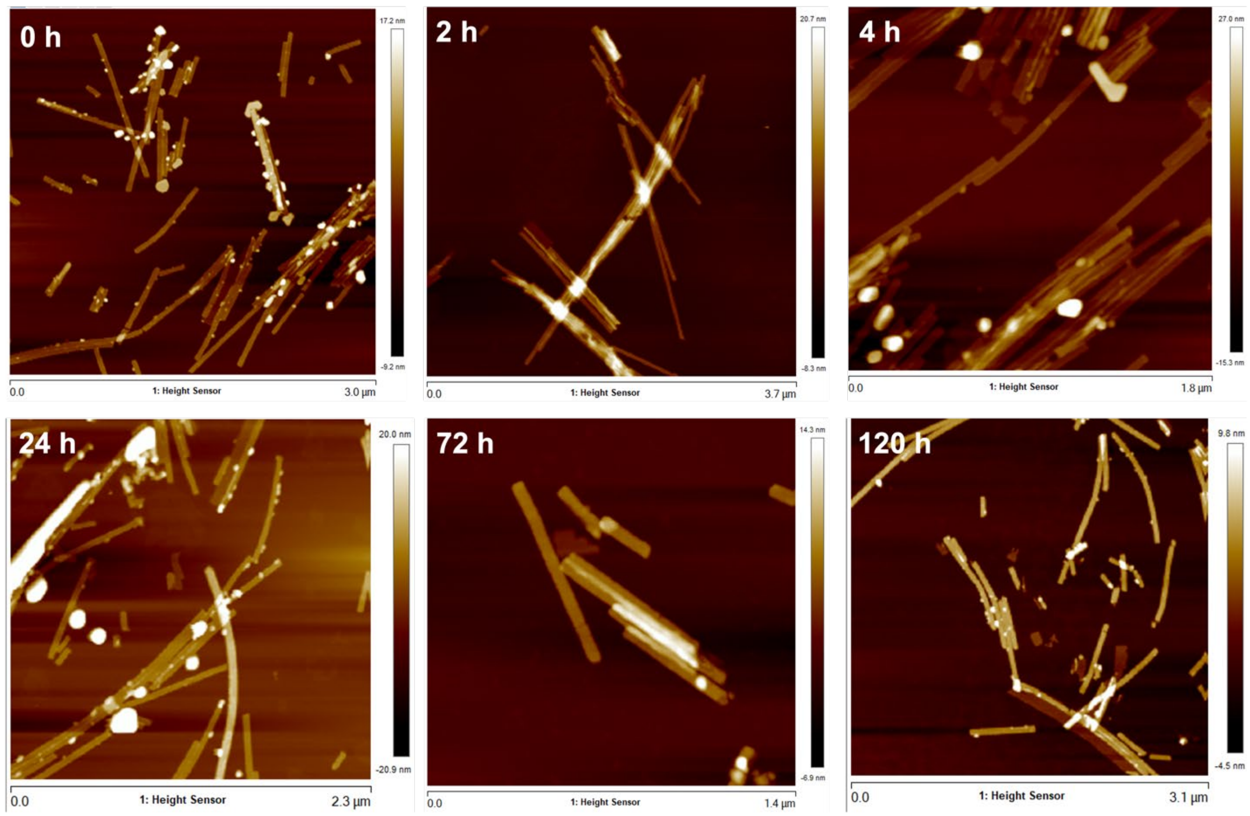


Figure S6. AFM images showing the structure of NAM-PNTs in PBS for up to 120 hours.

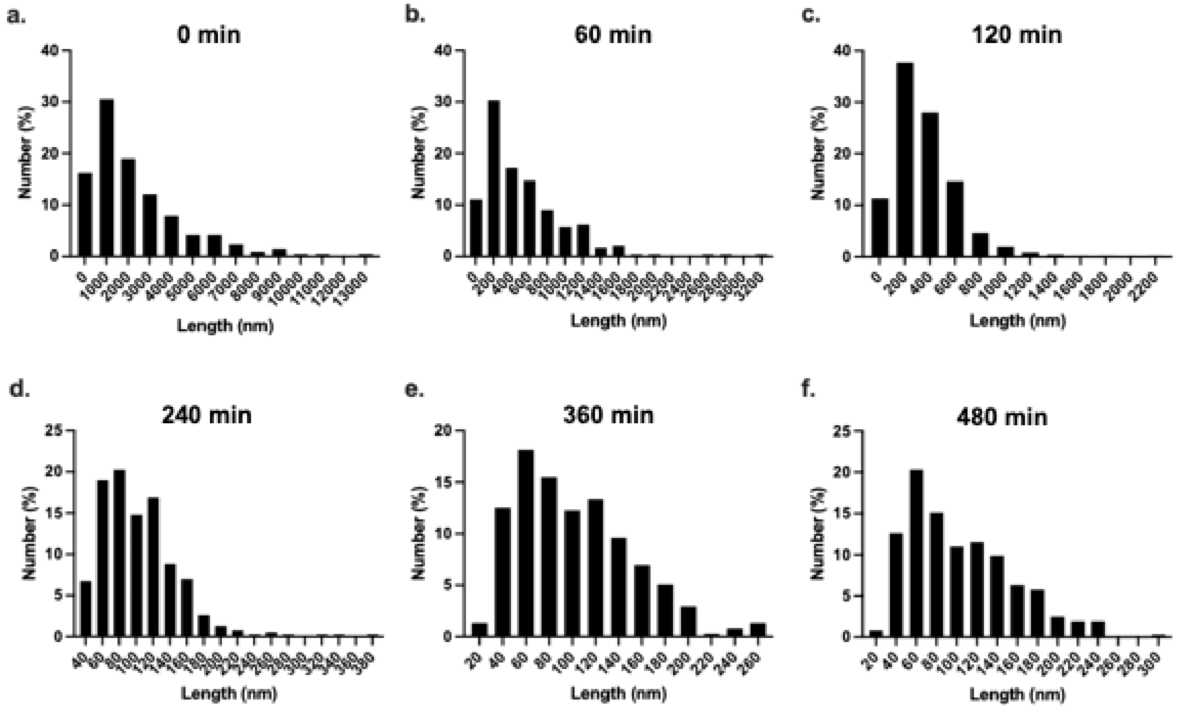


Figure S7. PNT length frequency distribution at different sonication times from 0 to 480 min.

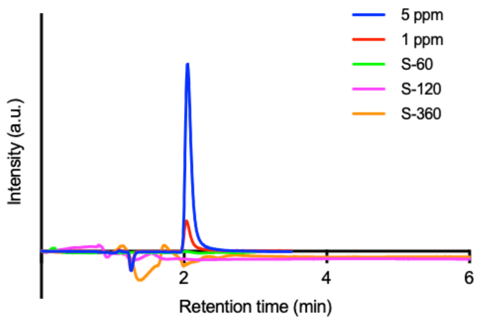


Figure S8. No NAM from NAM-PNT dissolved in the solution following up to 360 min of sonication. NAM was measured via liquid chromatography. 5 ppm and 1 ppm NAM solutions serve as standards for comparison.

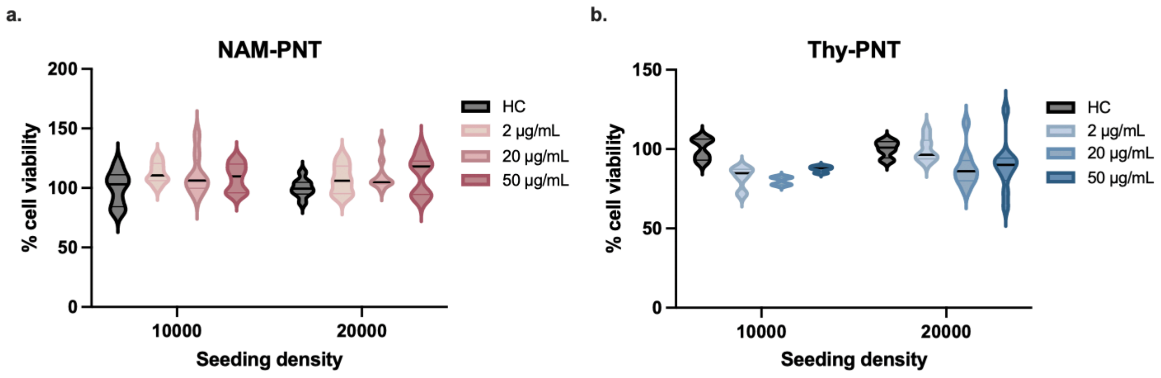


Figure S9. Seeding population effect on cell viability for a) NAM-PNT and b) Thy-PNT. Data are presented as violin plots that show the full range with lines at the median with interquartile range. (n=8-10).

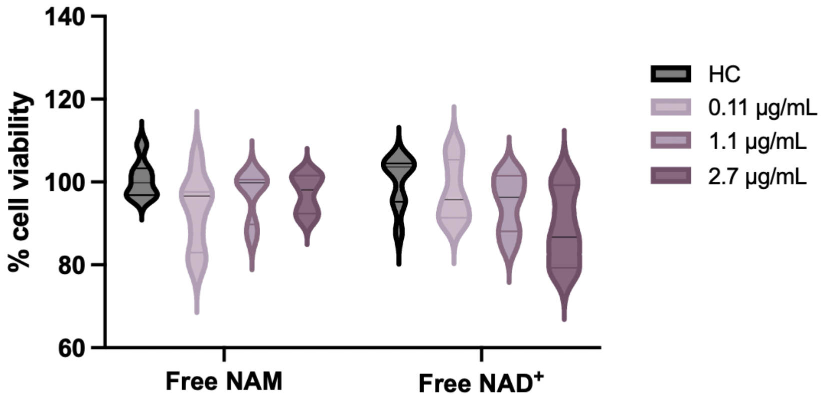


Figure S10. MTT data showing healthy BV-2 cells (HC) treated with 3 doses of free NAM or free NAD+ for 24 h. Data are presented as violin plots that show the full range with lines at the median with interquartile range. (n=7-8).

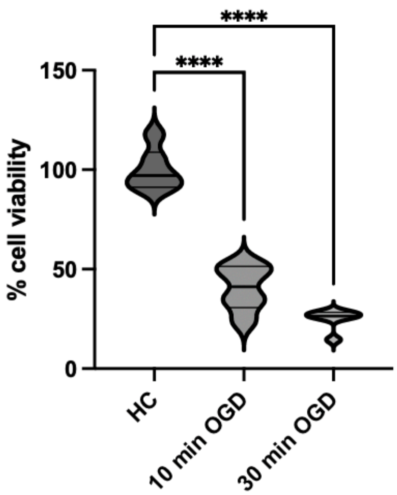


Figure S11. Difference in cell viability, as measured by MTT assay, after 10 min and 30 min of OGD compared to the healthy control (HC). Statistical significance was calculated via a Kruskal-Wallis test. Data are presented as violin plots that show the full range with lines at the median with interquartile range. (n = 6).

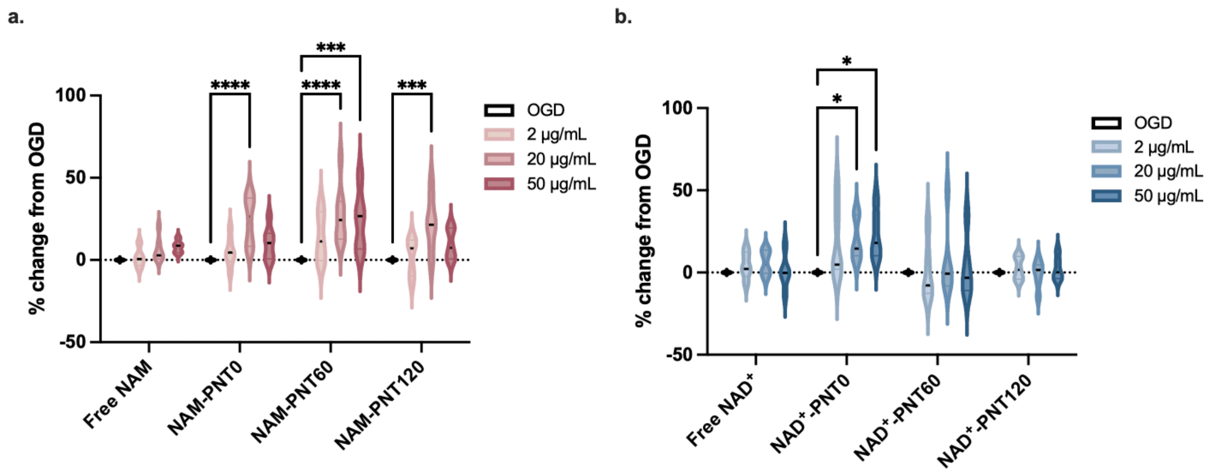


Figure S12. Change in metabolic activity of a) free NAM, and three lengths of NAM-PNTs, and b) free NAD+ and three lengths of NAD+-PNTs compared to the non-treated OGD cells. A dose-dependent effect was also observed, where 20 µg/mL NAM-PNTs led to the highest % change from OGD. Statistical significance was calculated via ordinary two-way ANOVA. Data are presented as violin plots that show the full range with lines at the median with interquartile range. (n=5-8).

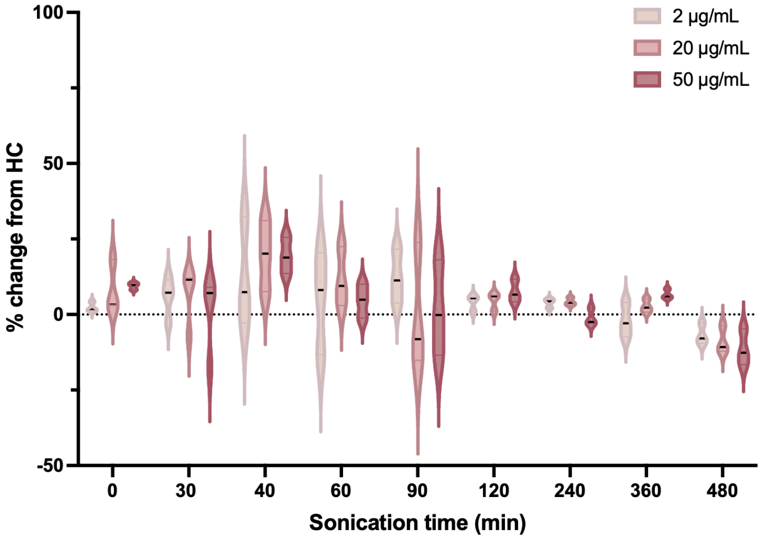


Figure S13. Change in metabolic activity of healthy BV-2 cells with the addition of 3 different doses of PNTs of different lengths generated by different sonication times compared to HC. Data are presented as violin plots that show the full range with lines at the median with interquartile range. (n = 3-4).

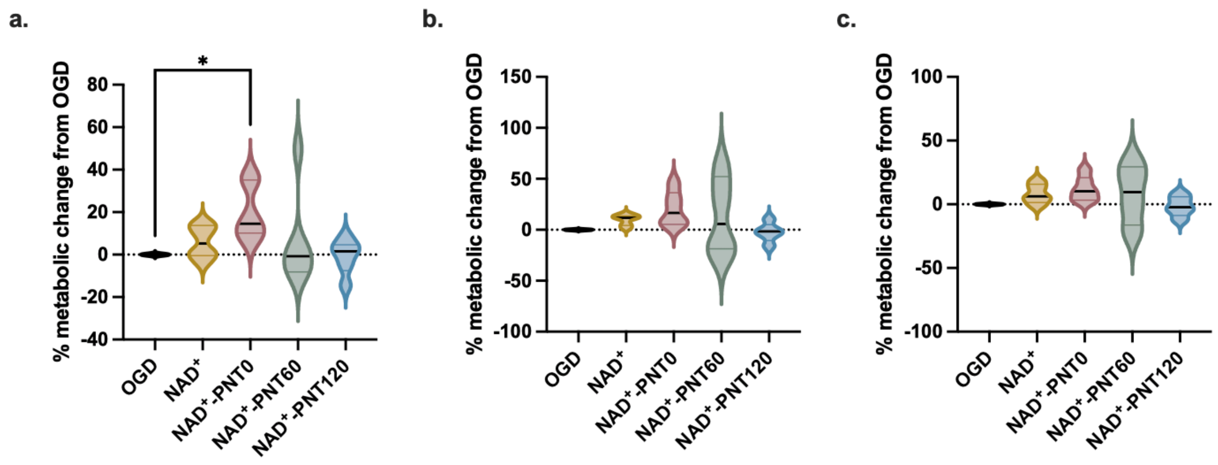


Figure S14. Sonication time impacts cell viability compared to non-treated OGD exposed controls for 20µg/mL of NAD+ delivered via PNTs with a) 24 h, b) 48 h and c) 72 h. Statistical significance was calculated via a Kruskal-Wallis test. Data are presented as violin plots that show the full range with lines at the median with interquartile range. (n=4-6).

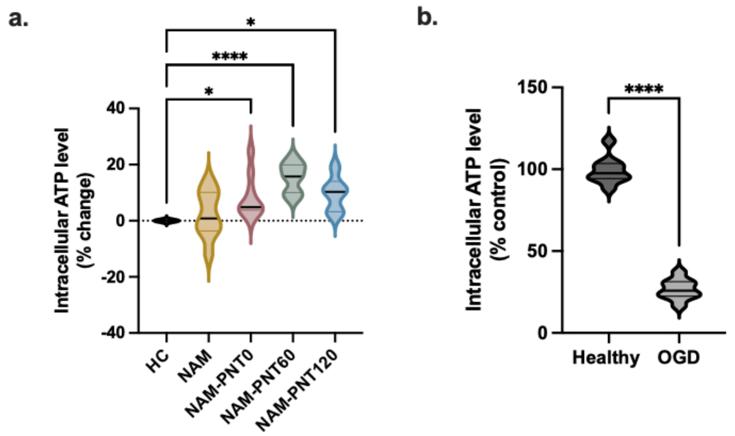


Figure S15. Intracellular ATP. a) Healthy BV-2 cells treated with NAM-PNTs showed an increase in intracellular ATP. b) Intracellular ATP is decreased in response to OGD-exposed in BV-2 cells. Statistical significance was calculated via a Kruskal-Wallis test. Data are presented as violin plots that show the full range with lines at the median with interquartile range. (n = 6-9).

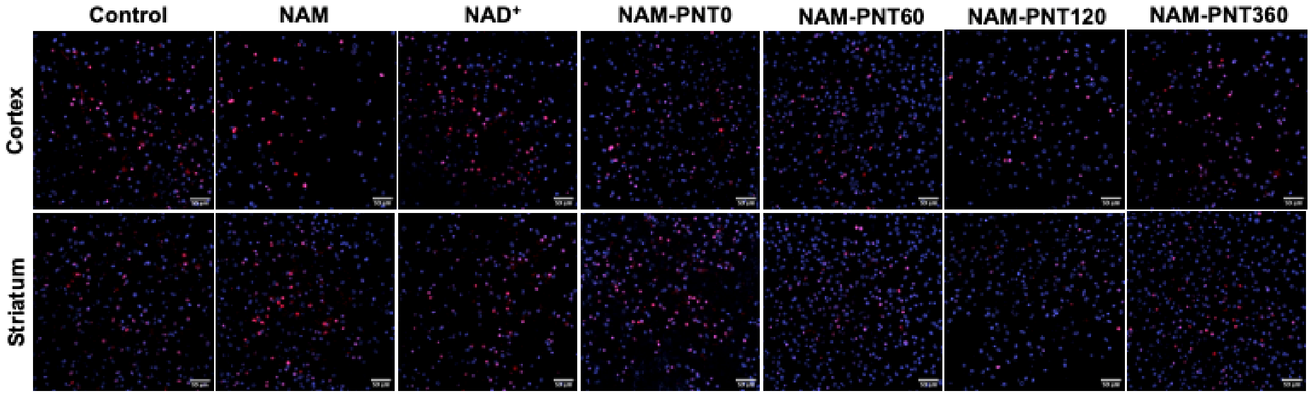


Figure S16. Representative images of propidium iodide signal (red) to visualize the amount of cell death after OGD or with 24 h of NAM, NAD, or NAM-PNT treatment applied immediately after OGD. Nuclei are stained with DAPI (blue). Scale bar, 50 µm.

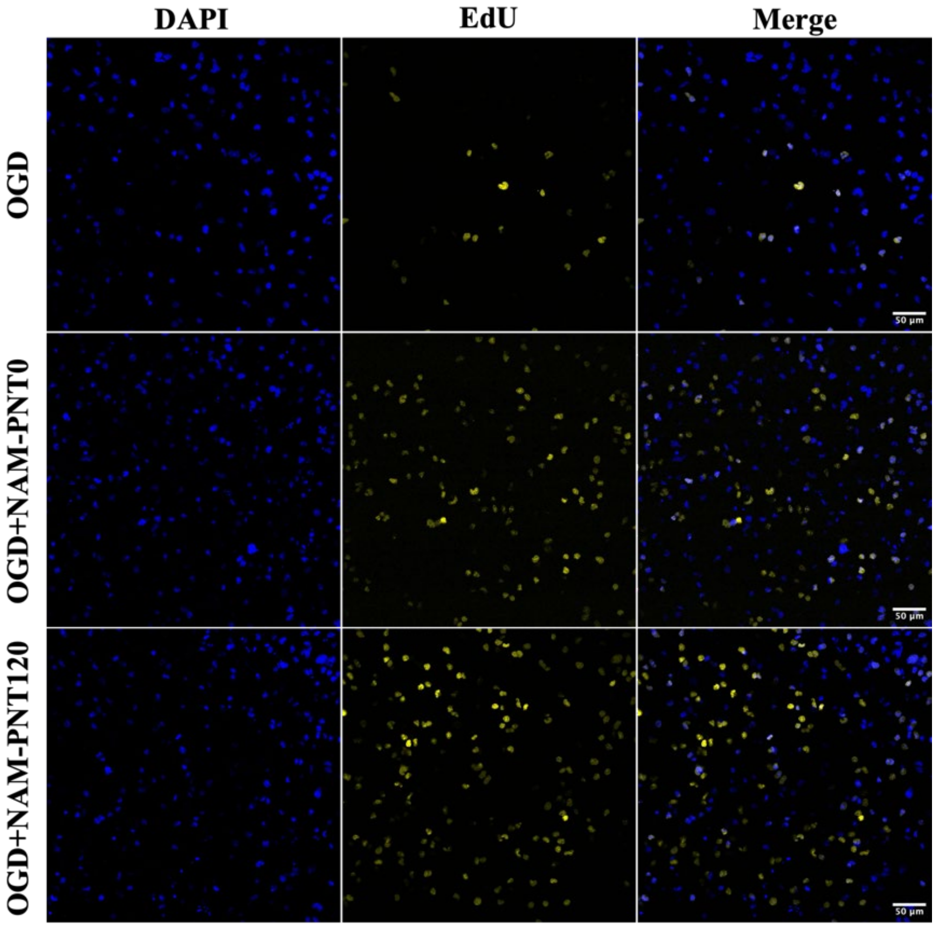


Figure S17. Representative images of EdU+ cells (yellow) 24 h after 30 min OGD at 4DIV from P10 slices. Nuclei are stained with DAPI (blue). Scale bar, 50 μm.

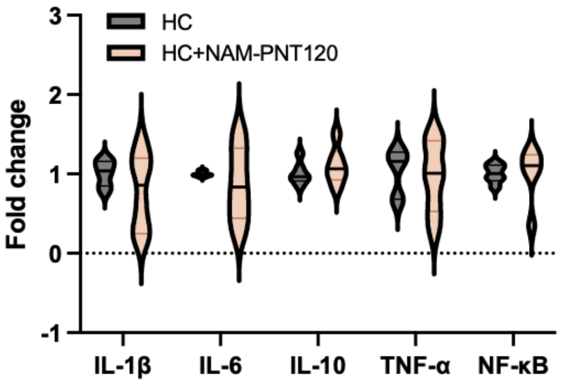


Figure S18. Fold-changes of mRNA markers for healthy slices with and without NAM-PNT treatment. Data are presented as violin plots that show the full range with lines at the median with interquartile range.

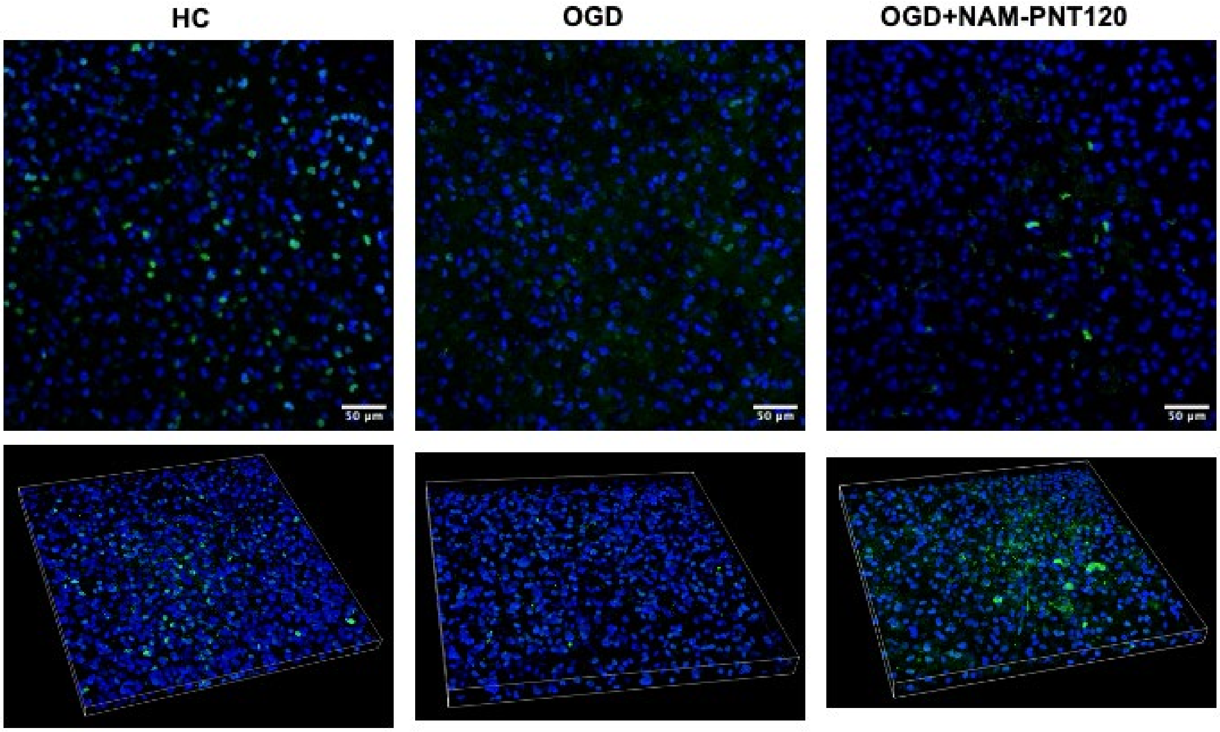


Figure S19. PAR signal (green) shows the extent of DNA damage in healthy, OGD, and OGD with NAM-PNT120 OWH brain slices. Nuclei are stained with DAPI. Scale bar, 50 µm.

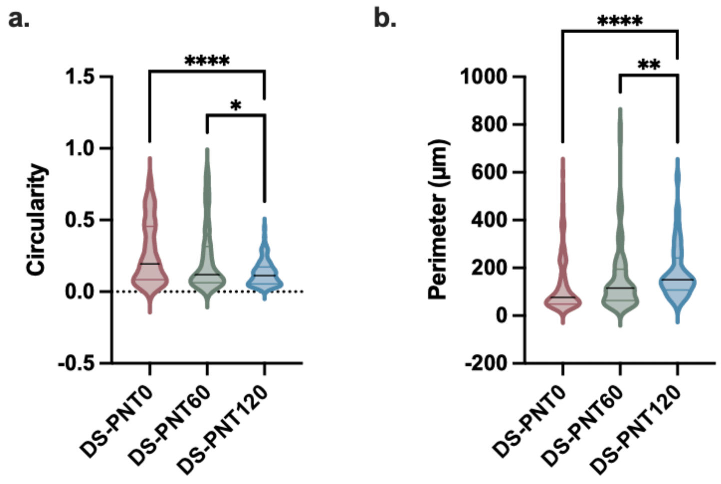


Figure S20. Microglia morphological analysis of a) circularity and b) perimeter in healthy OWH slices after exposure to DS-PN0, DS-PNT60 and DS-PNT120 for 24 h. Statistical significance was calculated via a Kruskal-Wallis test. Data are presented as violin plots that show the full range with lines at the median with interquartile range.

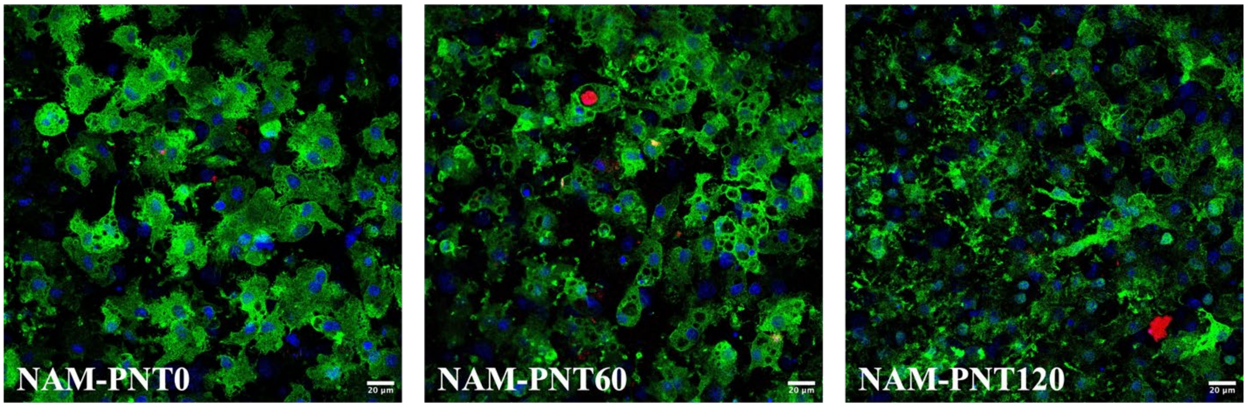


Figure S21. DS-PNT (red) localization in Iba1+ microglia (green) and ToPro-3 stained nuclei (blue) in OGD-exposed slices. Scale bar, 20 µm.

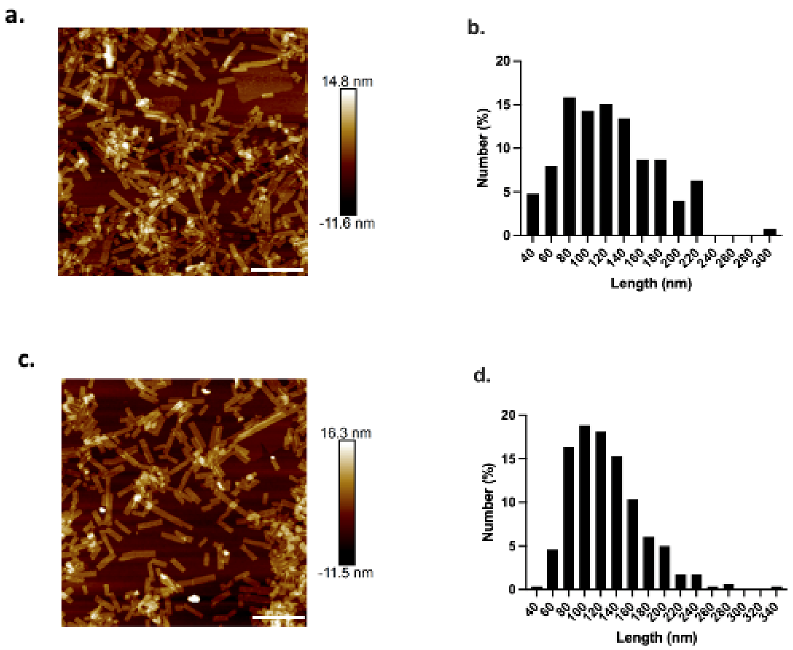


Figure S22. Representative AFM images and size distribution of a-b) PNT and c-d) NAM-PNT use for the in vivo study in HI injured rat pups. With 10 minutes of sonication-based emulsification, PNT has length of 124.24 ±49.83 nm and NAM-PNT of 128.16 ± 48.35 nm. Scale bar, 200 nm.

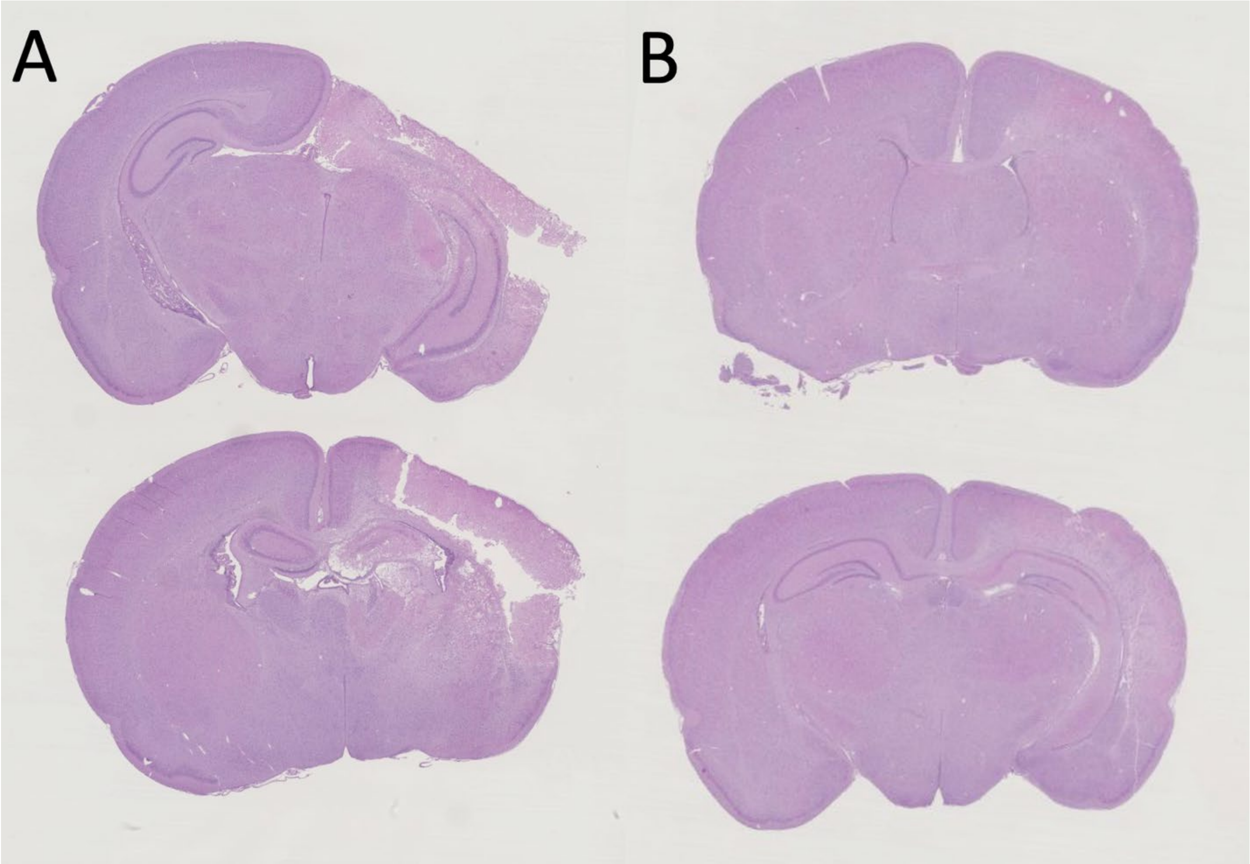


Figure S23. Representative images from a) Vehicle and b) NAM-PNT pups exposed to HI at P10. Increased percent area loss was observed in the vehicle group compared to the NAM-PNT group. Brains are from the media animal for each experimental group and are both female brains.

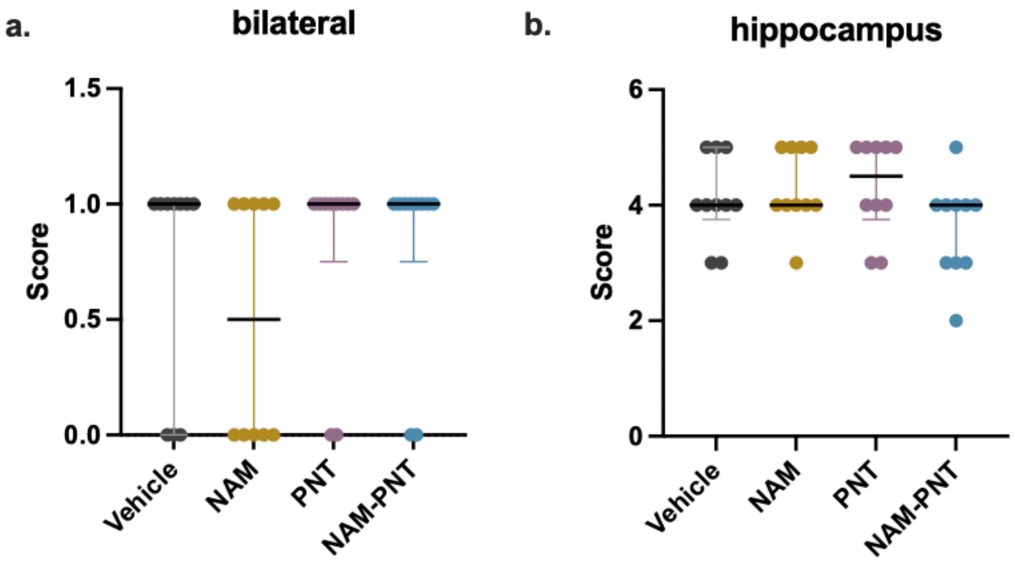


Figure S24. a) bilateral injury from pathology (score of 1) or unilateral injury (score of 0). b) hippocampus neuropathology data score of 0-5. Data are presented as scatter dot plots with lines at the median with interquartile range.

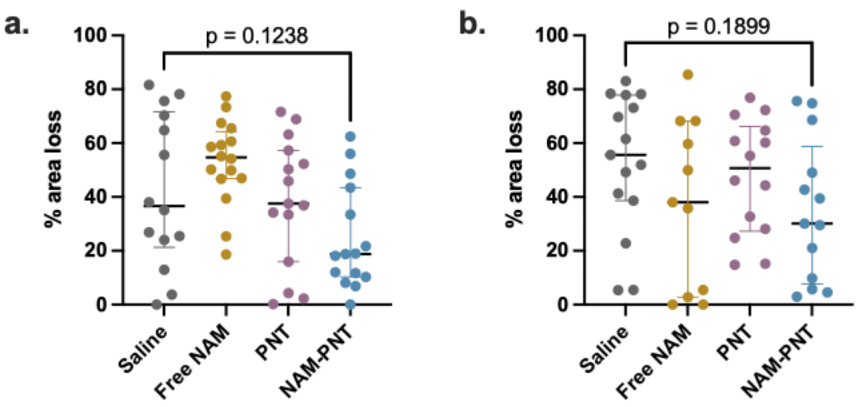


Figure S25. Total area of loss of a) male and b) female were calculated by assessing the percent area of tissue lost in the ipsilateral hemisphere, normalized to the contralateral hemisphere. Statistical significance was calculated via a Kruskal-Wallis test. Data are presented as scatter dot plots with lines at the median with interquartile range.

Table S1. Average NAM-PNTs length and aspect ratio under different sonication times.

| | 0 min | 30 min | 60 min | 120 min | 240 min | 360 min | 480 min |
|---|---|---|---|---|---|---|---|
| Ave length (nm) | 2343.61± 2249.96 | 561.29± 699.26 | 534.00± 479.58 | 360.78± 261.05 | 103.60± 47.45 | 102.13± 49.80 | 102.93 ± 51.41 |
| Aspect ratio | 332.90 | 79.73 | 75.85 | 51.25 | 14.72 | 14.51 | 14.62 |
